# Disordered C-terminal domain drives spatiotemporal confinement of RNAPII to enhance search for chromatin targets

**DOI:** 10.1101/2023.07.31.551302

**Authors:** Yick Hin Ling, Ziyang Ye, Chloe Liang, Chuofan Yu, Giho Park, Jeffry L. Corden, Carl Wu

## Abstract

Efficient gene expression requires RNA Polymerase II (RNAPII) to find chromatin targets precisely in space and time. How RNAPII manages this complex diffusive search in 3D nuclear space remains largely unknown. The disordered carboxy-terminal domain (CTD) of RNAPII, which is essential for recruiting transcription-associated proteins, forms phase-separated droplets *in vitro*, hinting at a potential role in modulating RNAPII dynamics. Here, we use single-molecule tracking and spatiotemporal mapping in living yeast to show that the CTD is required for confining RNAPII diffusion within a subnuclear region enriched for active genes, but without apparent phase separation into condensates. Both Mediator and global chromatin organization are required for sustaining RNAPII confinement. Remarkably, truncating the CTD disrupts RNAPII spatial confinement, prolongs target search, diminishes chromatin binding, impairs pre-initiation complex formation, and reduces transcription bursting. This study illuminates the pivotal role of the CTD in driving spatiotemporal confinement of RNAPII for efficient gene expression.

## Introduction

The carboxy-terminal domain (CTD) of RPB1 (Rpo21 in yeast), the largest subunit of RNA Polymerase II (RNAPII), is an essential and unique structural feature crucial for transcription regulation in eukaryotes. Defined by its intrinsically disordered structure^1^ and consensus heptad repeats (YSPTSPS)^2^, the CTD acts as a docking platform for recruiting transcription-associated proteins to interact with RNAPII^3–5^. Mutations or truncations of CTD repeats affect cell viability^6–10^ and impair gene activation^11–17^. Notably, the number of the heptad repeats in the CTD correlates with genome size throughout eukaryotes^13,18,19^.

The biochemical and biophysical properties of intrinsically disordered regions (IDRs), including the CTD, allow liquid-liquid phase separation (LLPS) through weak-multivalent transient interactions^20–22^. This results in the formation of phase-separated transcription condensates comprising RNAPII, the Mediator complex and other transcription-related proteins. These condensates have been proposed to play a critical role in organizing the transcription machinery to facilitate gene regulation^20–25^. By creating a specialized subnuclear environment, LLPS may help increase the concentration of RNAPII and associated factors for efficient transcription^26,27^. *In vitro*, the size of the phase-separated droplets formed by the biochemically purified CTD is dependent on its length. For example, the human RNAPII CTD, which has 52 heptad repeats, twice as many as its yeast counterpart with 26 repeats, readily forms larger droplets^20,22^. However, there is considerable debate concerning the relationship between *in vitro* reconstituted LLPS and its role as the primary mechanism in the formation of specific membraneless compartments within the cell^28–31^. Specifically, whether phase-separated RNAPII condensates represent a universally conserved mechanism, or if weak interactions driven by disordered CTDs can guide RNAPII diffusion without transitioning into a condensed phase in the living cell remains unclear. Here, we combined live-cell, single-molecule tracking with yeast genetics to elucidate the intricate diffusive mechanisms of RNAPII and the causal dependencies of its target search. Our results show that the freely diffusing RNAPII is spatially confined, but does not instigate formation of phase-separated condensates on a microscopic scale. The spatial confinement of RNAPII is governed by the full-length CTD, and requires both Mediator and global yeast chromatin organization. Importantly, such confinement allows freely diffusing RNAPII to stochastically oversample subnuclear regions enriched with active genes, establishing a highly efficient target search mechanism to enhance pre-initiation complex (PIC) formation and transcription bursting. Our work demonstrates how RNAPII leverages its search strategy as a rate-limiting step to promote transcription activation, providing insights on the functional significance of the CTD in regulating the spatiotemporal dynamics of RNAPII.

### Spatiotemporal mapping of transcription machinery

We performed live-cell single-molecule tracking (SMT) of fluorescently labeled endogenous RNAPII and other transcription proteins fused to HaloTag, and expressed as sole source under native promoter control (Fig. 1b, Extended Data Fig. 1e). Molecule trajectories were categorized into chromatin-bound or freely diffusing states based on diffusivity, geometry and angular orientation using a multi-parameter classification approach (Fig. 1a, Extended Data Fig. 1d and 2, and Supplementary Methods). To extract the spatiotemporal dynamics of the transcription machinery in the small budding yeast nucleus, we developed a mapping method for a strain with GFP markers delineating the plasma membrane, nuclear envelope and the nucleolus (Extended Data Fig. 1a-c and Supplementary Video 1). The method involves selection of G1 phase cells exhibiting minimal nuclear drift with distinct nuclear envelope and nucleolar boundaries (Extended Data Fig. 1g and 3a), normalization of trajectory positions based on nuclear landmarks, and superposition of single-molecule localizations from multiple (>10) cell nuclei on a 2D plane (Fig. 1a). This maps the spatial distribution of chromatin-bound and freely diffusing proteins over an integrated timeframe, and is grounded in the physical constraints imposed on the centromeres and telomeres of yeast interphase chromatin^32–35^ (Extended Data Fig. 1b). A similar mapping approach was previously described for the spatial probability density of single gene loci in yeast^36^. Likewise, we interpret the presented spatiotemporal map density as the probability density of protein localization over time, rather than instantaneous protein clustering.

**Fig. 1.**
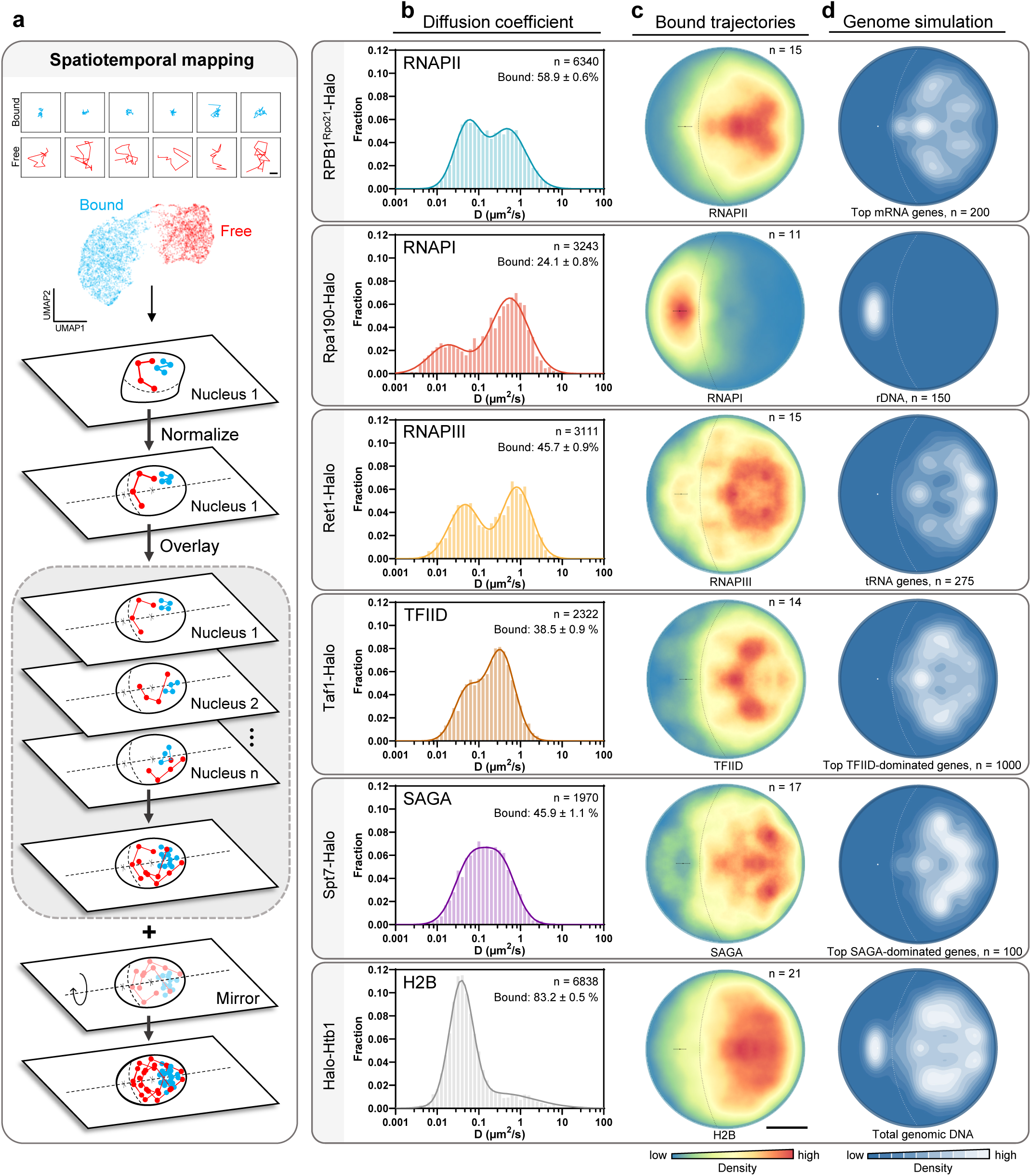
Spatiotemporal mapping of transcription machinery in living yeast nucleus. **a**, Spatiotemporal mapping workflow: bound and free trajectories (Scale bar: 0.2 µm) classified by multi-parameter analysis, and single-molecule localization and nuclear radius normalized to a circular model of 1 µm radius. Localizations overlaid from at least 10 nuclei that display minimal drift during image acquisition. Visualization of the spatiotemporal map on a single plane facilitated by mirroring around the central axis^36^. **b**, Diffusion coefficient histograms for RNAPII, RNAPI, RNAPIII, TFIID, SAGA and H2B. Representative HaloTag fusions shown (n: number of trajectories; mean value ± s.d.). **c**, Detection map of chromatin-bound trajectories. Scale bar: 0.5 µm (Error bar: centroid of nucleolus ± s.d.; n: number of nuclei). **d**, Simulation of the spatial distribution of gene classes in yeast nucleus (n: number of genes).

We validated the method by mapping the trajectories of free HaloTag, chromatin-bound centromeric Cse4 and telomeric Sir4 to the expected centromere (Cse4) and nuclear periphery regions (Sir4) established in the literature^32–35^ (Extended Data Fig. 3f,g). We next examined essential transcription proteins RNAPII, RNAPI, RNAPIII, TFIID, and SAGA, showing their free and bound diffusion coefficient distributions and corresponding fractions (Fig. 1b), and the spatiotemporal distribution of bound populations (Fig. 1c). As a reference for the distribution of bound transcription proteins, we generated a 2D projection of a 3D yeast genome model derived from 3C experimental data^37^ (Extended Data Fig. 3d). Integrating sequence coordinates (obtained from Saccharomyces Genome Database) of highly expressed genes across multiple gene classes^38^ allowed us to highlight their probability density distributions on the genome simulation map. Other investigations have similarly utilized 3D yeast genome models derived from polymer chain simulation to determine the probability densities of specific gene classes, such as tRNA genes and highly expressed mRNA genes^39,40^. Unlike the semi-homogenous H2B density map (Fig. 1c; bottom), the bound transcription protein maps (Fig. 1c) are uniquely distinct, and spatially show resemblance to the corresponding functional gene classes located by genome simulation^37^ (Fig. 1d). Notably, chromatin-bound TFIID and SAGA exhibit distinct spatial territories in the spatiotemporal maps (Fig. 1c), mirroring the TATA-less (TFIID-dominated) and TATA-containing (SAGA-dominated) genes^41^ respectively in the genome simulations (Fig. 1d), raising the possibility that the regulation of gene classes may be related to their spatial positioning^39,40^. Further investigation is needed to fully understand the implications of this organization.

### CTD governs chromatin binding and spatiotemporal confinement of free RNAPII

Given structural similarities between the three eukaryotic RNA polymerases, we compared the spatiotemporal distribution and diffusivity of RNAPII with RNAPI and RNAPIII. Freely diffusing RNAPII shows a higher degree of nucleolar exclusion (Extended Data Fig. 4a-c), and diffuses more slowly than expected from its molecular weight (Extended Data Fig. 4d). We hypothesized that the distinct spatiotemporal dynamics of freely diffusing RNAPII might be partially attributed to its large hydrodynamic radius due to the presence of the disordered CTD on its largest subunit, RPB1^Rpo21^ ^42,43^, a feature absent from RNAPI and RNAPIII (Fig. 2a and Extended Data Fig. 4e,f). Notably, we observed that increasing the number of CTD heptad repeats across a spectrum of 0, 8, 9, 10, 15, 20, 26 (wildtype), 52, 78, to 104 repeats led to increased nucleolar exclusion and decreased diffusion coefficient of free RNAPII (D_free_) (Fig. 2b and Extended Data Fig. 4g). This indicates that CTD length might directly and/or indirectly modulate the diffusivity and distribution of RNAPII between nucleolus and nucleoplasm. A large hydrodynamic radius, conferred by a long and highly disordered CTD, would slow RNAPII and limit diffusion into the nucleolus via size exclusion, thereby increasing its effective concentration in the nucleoplasm. As noted below, there are additional mechanisms relevant to RNAPII nucleoplasmic distribution.

**Fig. 2.**
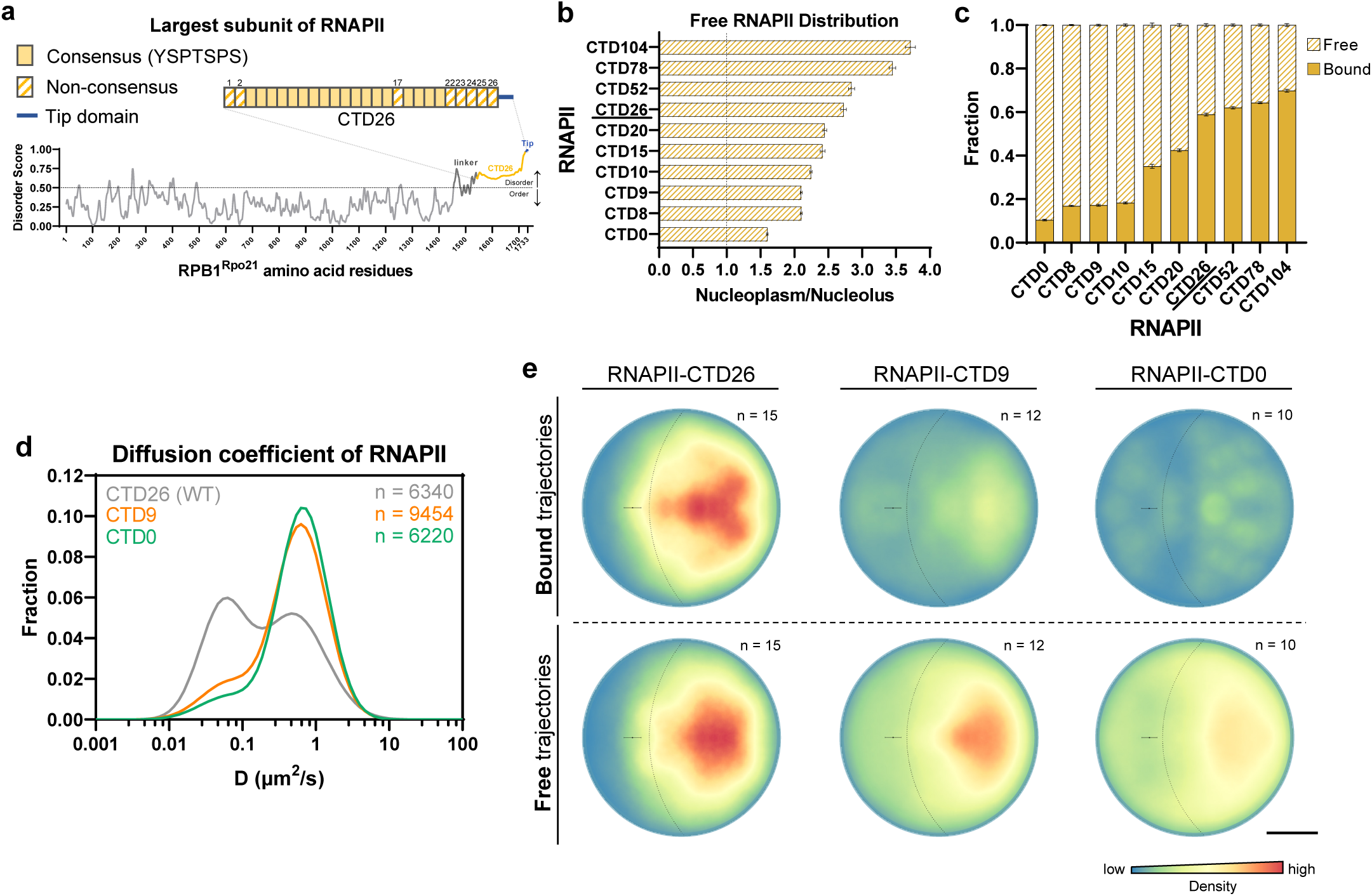
CTD governs spatiotemporal dynamics of RNAPII. **a**, Disordered C-terminal domain of RPB1^Rpo21^ consisting of 26 heptad repeats with consensus sequence YSPTSPS. IUPRED3 scores above 0.5 considered disordered. **b**-**e**, Spatiotemporal dynamics of RNAPII and CTD mutants characterized by **b**, ratio of freely diffusing molecules between the nucleoplasm and nucleolus (n = 100 resamplings; mean value ± s.d.), **c**, fraction of free and bound (n = 100 resamplings; mean value ± s.d.), **d**, two-component Gaussian fit of diffusion coefficient histograms, and **e**, spatiotemporal mapping for bound and free trajectories. Scale bar: 0.5 µm (Error bar: centroid of nucleolus ± s.d.; n: number of nuclei). Top left panel is reuse of Fig. 1c top panel.

Next, we explored the impact of CTD length on the chromatin-bound population of RNAPII. Progressive truncation of the CTD from 26 repeats to 20, 15, 10, 9, 8, and 0 repeats notably reduces its bound fraction, whose low diffusion coefficient is comparable to bound H2B (D_bound_ ∼0.05 µm^2^/s) (Fig. 1b, 2d, Extended Data Fig. 5a and Supplementary Video 2). Conversely, extending the CTD to 52, 78, and 104 repeats shows a trend toward an increase in this fraction, suggesting potential enhancement of chromatin association (Fig. 2c and Extended Data Fig. 5a). This trend remains consistent after normalization of nuclear RPB1^Rpo21^ fluorescence to the expression level across mutants^13^ (Extended Data Fig. 5c,d). We primarily focus on CTD truncations due to their pronounced effects; The spatiotemporal maps show a global reduction of RNAPII binding for truncated CTD strains CTD9 and CTD0 (Fig. 2e; top). Importantly, the map of free RNAPII shows spatial confinement in a subnuclear region enriched for the bound population (Fig. 2e), an area also abundant in highly expressed yeast genes (Fig. 1d and Extended Data Fig. 3e). This confinement is disrupted in the CTD9 mutant, and further reduced in CTD0 (Fig. 2e; bottom). Hence, the CTD has a critical role in the spatiotemporal dynamics for both bound and free RNAPII.

### RNAPII confinement characterized by anisotropic diffusion

We quantified the spatiotemporal confinement of freely diffusing RNAPII by measuring anisotropy, using the ratio of extreme backward to forward angles (f_180/0_) of free trajectories in the nucleoplasm, with confined trajectories showing backward bias^44^ (Fig. 3a,b; top and Supplementary Video 3). We observed a substantial backward bias in wildtype (WT) RNAPII-CTD26, which is largely absent for RNAPII-CTD9 and RNAPII-CTD0 mutants, and free HaloTag (Fig. 3a,b). On the other hand, CTDs expanded to 52, 78, and 104 repeats show increasing anisotropy (Extended Data Fig. 6c). A f_180/0_ peak around 150 nm (Fig. 3b) suggests RNAPII confinement within potential domains, as reported for CTCF in mammalian cells, in which anisotropic diffusion through transient trapping in zones’ has been proposed as an explanatory model^44^. Given that yeast interphase chromosomes are physically tethered to the nuclear envelope via telomeres, we disrupted global chromatin organization by inducing telomere detachment through deletion of *esc1* and *yku70*^45^ (Fig. 3c,d and Supplementary Video 4). This resulted in a slight reduction of RNAPII binding (Fig. 3e). Notably, telomere detachment abolishes the f_180/0_ peak at 150 nm, but maintains elevated f_180/0_ across length scales as compared to that of CTD truncation mutants (Fig. 3f,g). This suggests that global chromatin organization might be required for establishing transient trapping zones or domains^46^ that contribute to RNAPII confinement kinetics.

**Fig. 3.**
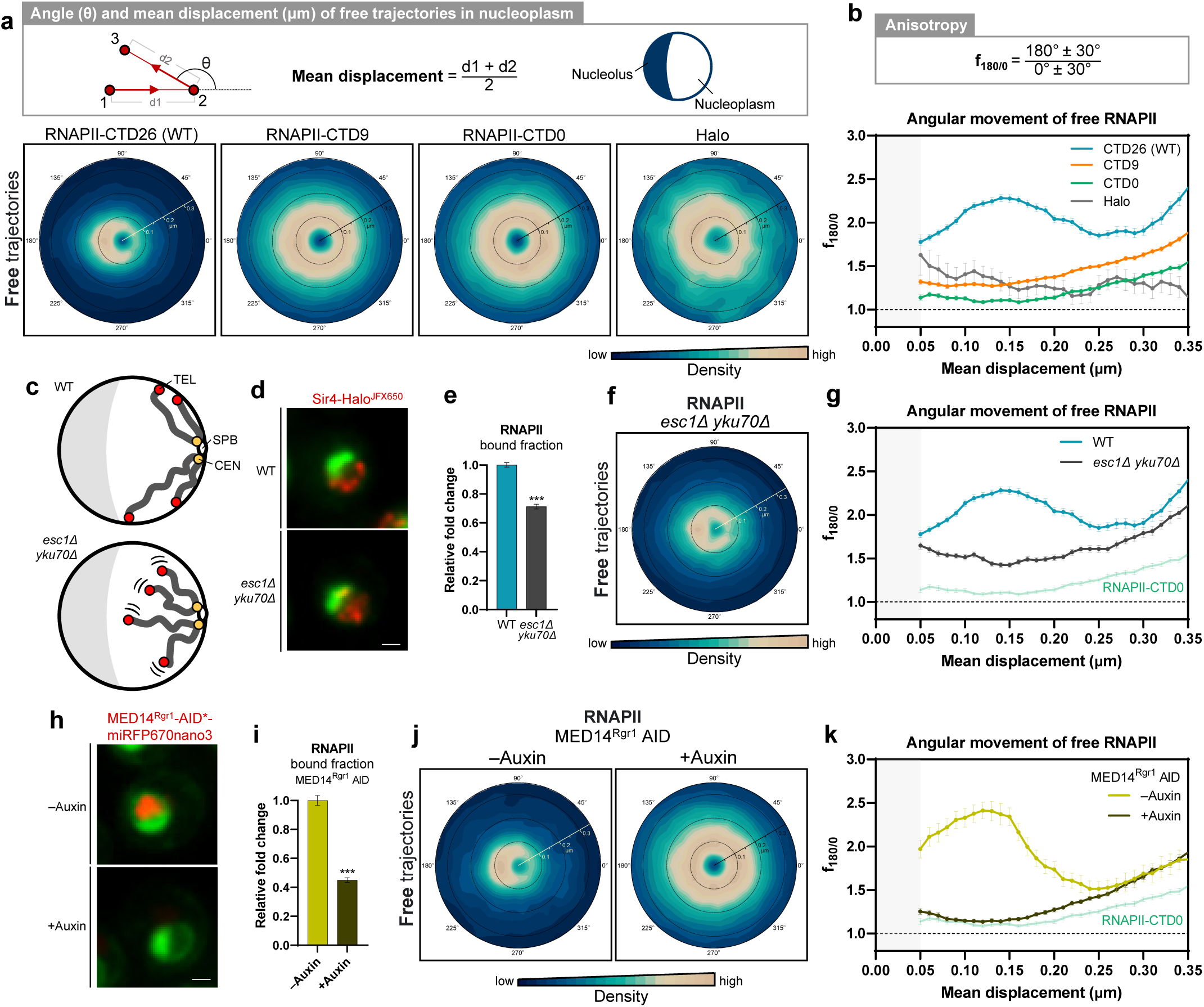
The interplay of CTD, chromatin organization and Mediator shapes RNAPII anisotropic diffusion. **a**, Angle (θ; angular axis) and mean displacement (radial axis; unit: µm) polar plots of RNAPII, CTD mutants and HaloTag (NLSx2-Halo). **b**, Anisotropic diffusion of freely diffusing of RNAPII, CTD mutants and HaloTag in nucleoplasm measured by f_180/0_ (n = 100 resamplings; mean value ± s.d.). **c**, Impaired telomere (TEL) attachment in *esc1*Δ *yku70*Δ strain. **d**, Fluorescence signal of Sir4-Halo^JFX650^ (red) in WT and *esc1*Δ *yku70*Δ. Nucleolar and ER markers in green. Scale bar: 1.0 µm (Supplementary Video 4). **e**, Relative bound fraction of RNAPII in WT and *esc1*Δ *yku70*Δ strains (n = 100 resamplings; mean value ± s.d.; two-tailed unpaired t-test). **f**, Angle and displacement polar plots of RNAPII in *esc1*Δ *yku70*Δ strain. **g**, f_180/0_ of RNAPII in WT and *esc1*Δ *yku70*Δ strains (n = 100 resamplings; mean value ± s.d.). **h**, Fluorescence signal of MED14^Rgr1^-AID*-miRFP670nano3 (red) before and after Mediator degradation by auxin treatment. Nucleolar and ER markers in green. Scale bar: 1.0 µm. **i**, Relative bound fraction of RNAPII after Mediator degradation (n = 100 resamplings; mean value ± s.d.; two-tailed unpaired t-test). **j**, Angle and displacement polar plots of RNAPII before and after Mediator degradation. **k**, f_180/0_ of RNAPII before and after Mediator degradation (n = 100 resamplings; mean value ± s.d.).

Given the reported weak interactions between mammalian Mediator IDRs and RNAPII CTD *in vitro*^22^, we investigated the role of Mediator in the diffusion confinement of free RNAPII. We observed a significant reduction of RNAPII binding when Mediator is degraded using an auxin-inducible degron strain (MED14^Rgr1^ AID) (Fig. 3h,i). Importantly, there is a large reduction of backward bias as shown in the angle distribution plots and anisotropy (f_180/0_) for free RNAPII (Fig. 3j,k). Furthermore, spatiotemporal mapping indicates a considerable decrease in free RNAPII confinement (and in the nucleoplasm/nucleolus ratio) following Mediator degradation (Extended Data Fig. 7b,d). For RNAPII-CTD9, Mediator degradation similarly decreased its confinement and further reduced the bound fraction (Extended Data Fig. 7j,k). Additionally, there is partial overlap between the probability density maps of bound Mediator and the confined trajectories of free RNAPII-CTD26 (Extended Data Fig. 7a). Collectively, we conclude that Mediator is required for RNAPII diffusion confinement in the nucleoplasmic compartment.

### No substantial RNAPII clustering observed in yeast nucleoplasmic compartment

The confined diffusion of RNAPII raises the question of phase separation in condensates. We did not observe fluorescent RNAPII puncta under wide-field microscopy in RPB1^Rpo21^-HaloTag fusions harboring WT or CTD mutants with reduced or expanded repeats (Fig. 4a). To investigate the possibility of nanoscale puncta, we performed fixed-cell stochastic optical reconstruction microscopy (STORM) on RPB1^Rpo21^-HaloTag co-stained with JF552 and JFX650. We imaged two color channels simultaneously and performed pairwise cross-correlation C(r) analysis to account for clustering artifacts due to fluorophore blinking^47^ (Fig. 4b). The C(r) values for RNAPII-CTD26, similar to those of H2B and RNAPII-CTD9, are close to 1, suggesting no significant clustering (Fig. 4c). A recent *in vitro* study suggests that multiple yeast RNAPII molecules could bind to a single UAS (upstream activating sequence)^48^. While our low C(r) values do not exclude the presence of minute or infrequent clustering, the bulk of RNAPII exhibits a dispersed distribution.

**Fig. 4.**
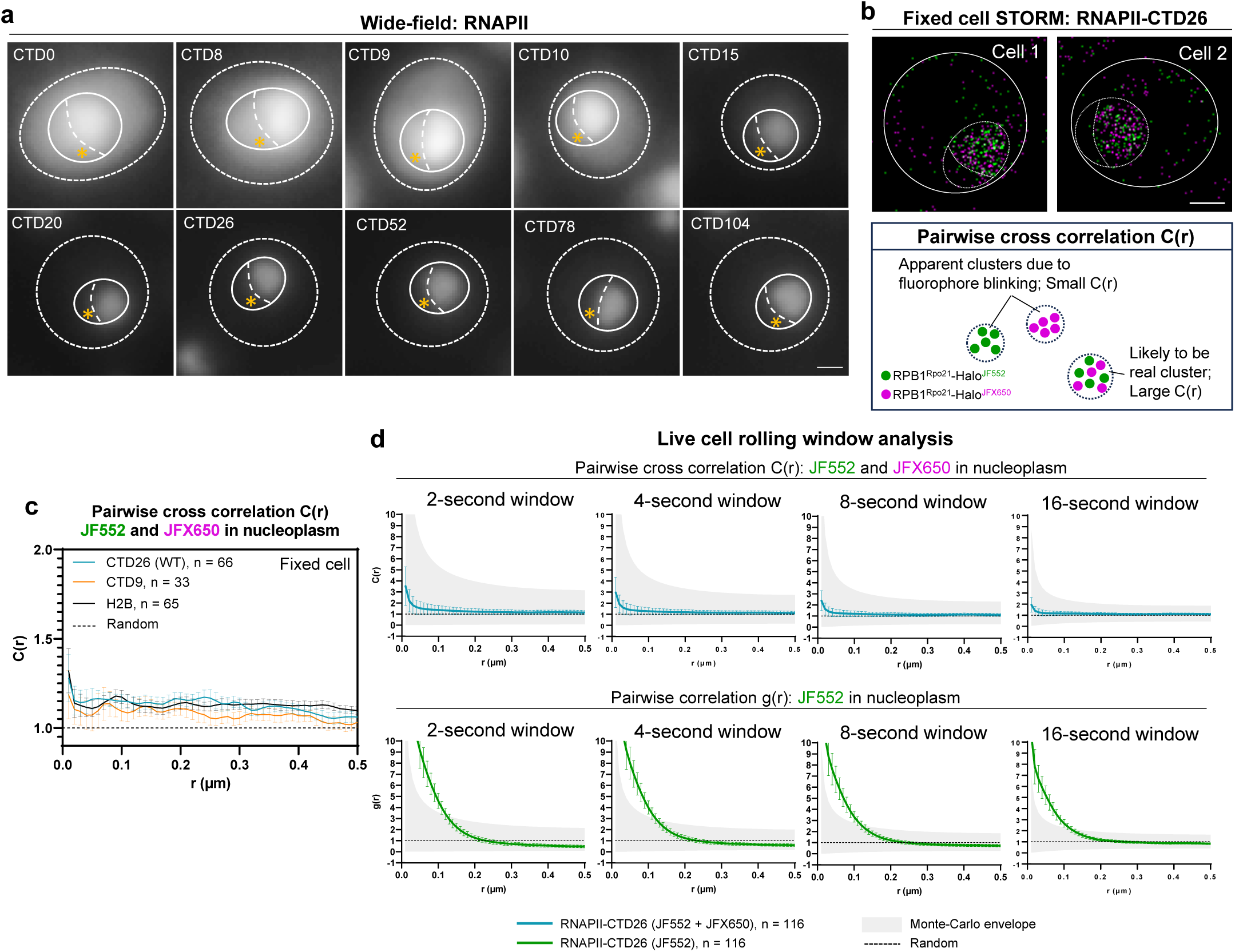
No substantial RNAPII clustering in yeast nucleoplasm. **a**, Bulk wide-field staining of RPB1^Rpo21^-Halo with JFX650 in live cells. Nucleolus indicated by asterisk. Scale bar: 1.0 µm. **b**, RPB1^Rpo21^-Halo fixed-cell STORM images using JF552 and JFX650. Scale bar: 1.0 µm (top). RNAPII clustering analyzed by pairwise cross-correlation C(r) (bottom). **c**, C(r) of RNAPII-CTD26, RNAPII-CTD9 and H2B in nucleoplasm (n: number of nuclei; mean value ± s.d.). **d**, RPB1^Rpo21^-Halo rolling window analysis for live cell two-color STORM (n: number of nuclei; mean value ± s.d.).

Given the possibility that transient clusters could be missed by imaging of fixed cells, we replicated the co-staining experiment using live yeast. We analyzed the data using multiple short time windows for potential time-correlated detections^24^, and performed Monte Carlo simulations to account for high C(r) values resulting from the limited number of detections within brief observation windows. Nonetheless, we found no strong statistical evidence for transient RNAPII clustering (Fig. 4d; top). Notably, pairwise correlation g(r) analysis performed using only JF552 indicated strong clustering due to repetitive fluorophore blinking or overcounting of fluorophores with extended lifetimes (Fig. 4d; bottom), which underscores the importance of two-color staining for reliable cluster assessment^47^. Thus, the observed high probability density of free RNAPII over time in our spatiotemporal map (Fig. 2e; bottom) results from transient confinement, rather than stable clustering or phase separation into condensates. This implies that despite the spatial confinement of RNAPII in subnuclear regions, its overall distribution at any given time remains broadly homogeneous. Consistent with this, Mediator degradation, which reduces RNAPII chromatin binding (Fig. 3i) and anisotropy (f_180/0_) (Fig. 3k), shows no substantial change in the diffusion coefficient of free RNAPII (Extended Data Fig. 7c). This suggests that the free RNAPII confinement is transient, and that the overall mobility over time, which is reflected in the diffusion coefficient, remains largely unchanged.

### CTD length is critical for efficient RNAPII target search kinetics

To further investigate the role of the CTD in controlling the chromatin binding kinetics of RNAPII, we measured the residence time of individual RNAPII binding events using the slow-tracking’ SMT mode that minimizes photobleaching with low laser power (Fig. 5b, Extended Data Fig. 8c,d and Supplementary Video 5). Halo-H2B cells were imaged as a spike-in control to correct for photobleaching and chromatin movements out of the focal plane (Fig. 5a and Extended Data Fig. 8f). We observed an average stable residence time of ∼20 s for WT RNAPII-CTD26, which encompasses nonproductive and productive pre-initiation, initiation, elongation, and termination. For RNAPII-CTD9, this average value is reduced substantially to ∼10 s, and the stably bound fraction decreases from ∼16% to ∼3% (Fig. 5c-e). Structural investigations have shown that CTD repeats make specific contacts with the Mediator complex^49^. The short residence time of RNAPII-CTD9 is therefore likely due to compromised pre-initiation complex (PIC) recruitment and engagement with the Mediator^50^ (Fig. 5j). Similarly, when Mediator is degraded, the WT RNAPII-CTD26 stably bound fraction is decreased, and the residence time even further reduced to ∼5 s (Fig. 5g-i), consistent with previous ChIP results showing depletion of RNAPII occupancy across nearly all genes under similar degron conditions^51^. Consistent with this, Mediator degradation also leads to an additional decrease in the stably bound fraction of RNAPII-CTD9, with residence time further reduced from ∼10 s to ∼4 s (Extended Data Fig. 7l-n).

**Fig. 5.**
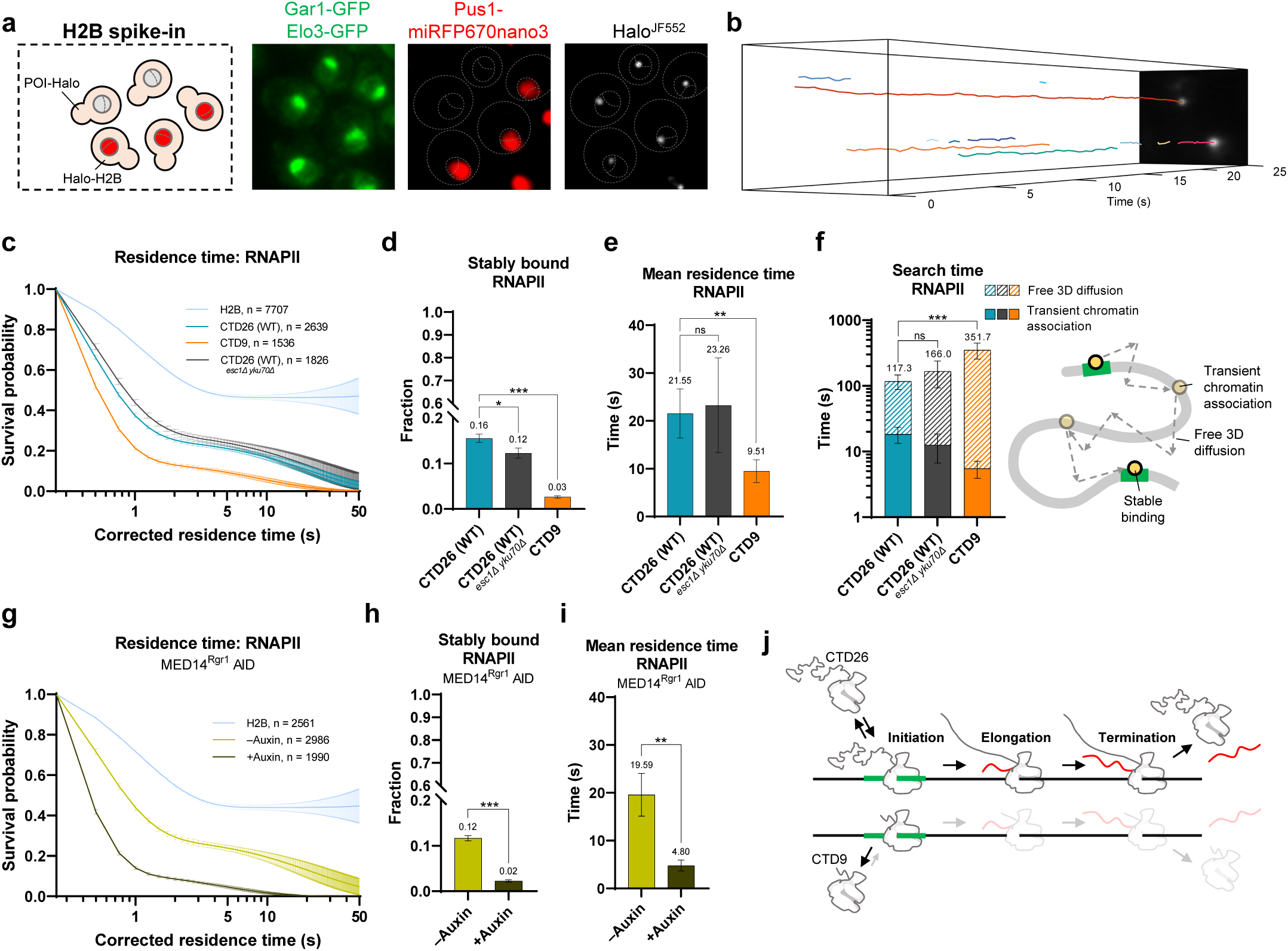
CTD facilitates RNAPII target-search kinetics. **a**, Halo-H2B cell spike-in for residence time correction with Pus1-miRFP670nano3 nuclear marker. Scale bar: 1.0 µm. **b**, Kymograph of slow tracking trajectories of RNAPII (RPB1^Rpo21^-Halo) (Supplementary Video 5). **c**, Survival probability of H2B-corrected residence times for bound RNAPII in CTD26 (WT), CTD9 and *esc1*Δ *yku70*Δ strains (n: number of trajectories; mean value ± s.d.). **d**-**f**, **d**, Stably bound fractions, **e**, mean residence times, and **f**, search times (left) of RNAPII in CTD26 (WT), CTD9 and *esc1*Δ *yku70*Δ strains (n = 10,000 resamplings; mean value ± s.d.; two-tailed unpaired t-test). **f**, Simplified mode of target search in the nucleus (right). **g**, Survival probability of H2B-corrected residence times of RNAPII before and after Mediator degradation (MED14^Rgr1^ AID) (n: number of trajectories; mean value ± s.d.). **h**-**i**, **h**, Stably bound fractions and **i**, mean residence times of RNAPII before and after Mediator degradation (n = 10,000 resamplings; mean value ± s.d.; two-tailed unpaired t-test). **j**, Model of reduction of RNAPII residence time in CTD9 mutant.

We estimated the target search time of RNAPII, i.e. the average search duration for an RNAPII molecule between two stable binding events including intervening 3D diffusion and multiple transient chromatin associations, to be ∼120 s (Fig. 5f). Although there is no significant change in target search time for the telomere attachment mutant *esc1*Δ *yku70*Δ, we found that the CTD9 mutant indeed shows prolonged RNAPII target search, increasing from the WT value of ∼120 s to ∼350 s (Fig. 5f). This observation is consistent with the marked reduction in spatiotemporal confinement of CTD9 and CTD0 mutants (Fig. 2e; bottom and 3b), suggesting that this confinement facilitates the RNAPII search for and association with promoter chromatin.

### CTD regulates chromatin-binding of PIC components and *HSP82* transcription bursting

In addition to facilitating RNAPII search for chromatin targets, CTD is a versatile scaffold coordinating the recruitment and release of transcription initiation, elongation and termination factors throughout the transcription cycle^3,4^. Accordingly, we examined the effects of CTD truncation on the dynamics of PIC components. In the CTD9 mutant, the bound fractions of TFIIA (Halo-Toa1), TFIIB (Sua7-Halo), TFIIE (Tfa1-Halo), TFIIF (Tfg1-Halo), and TFIIH (Tfb4-Halo and Kin28-Halo), but not TBP (Halo-Spt15), TFIID (Taf1-Halo), SAGA (Spt7-Halo and Spt8-Halo) and Mediator (Rgr1-Halo and Med1-Halo), are substantially decreased (Fig. 6a,b). This reduction is consistent with the crucial role of the CTD in maintaining the integrity of the PIC by facilitating key interactions^49,50,52^. The modest binding changes for two core subunits of Mediator (Rgr1 and Med1) suggest that other interactions besides CTD binding contribute to its stability on chromatin.

**Fig. 6.**
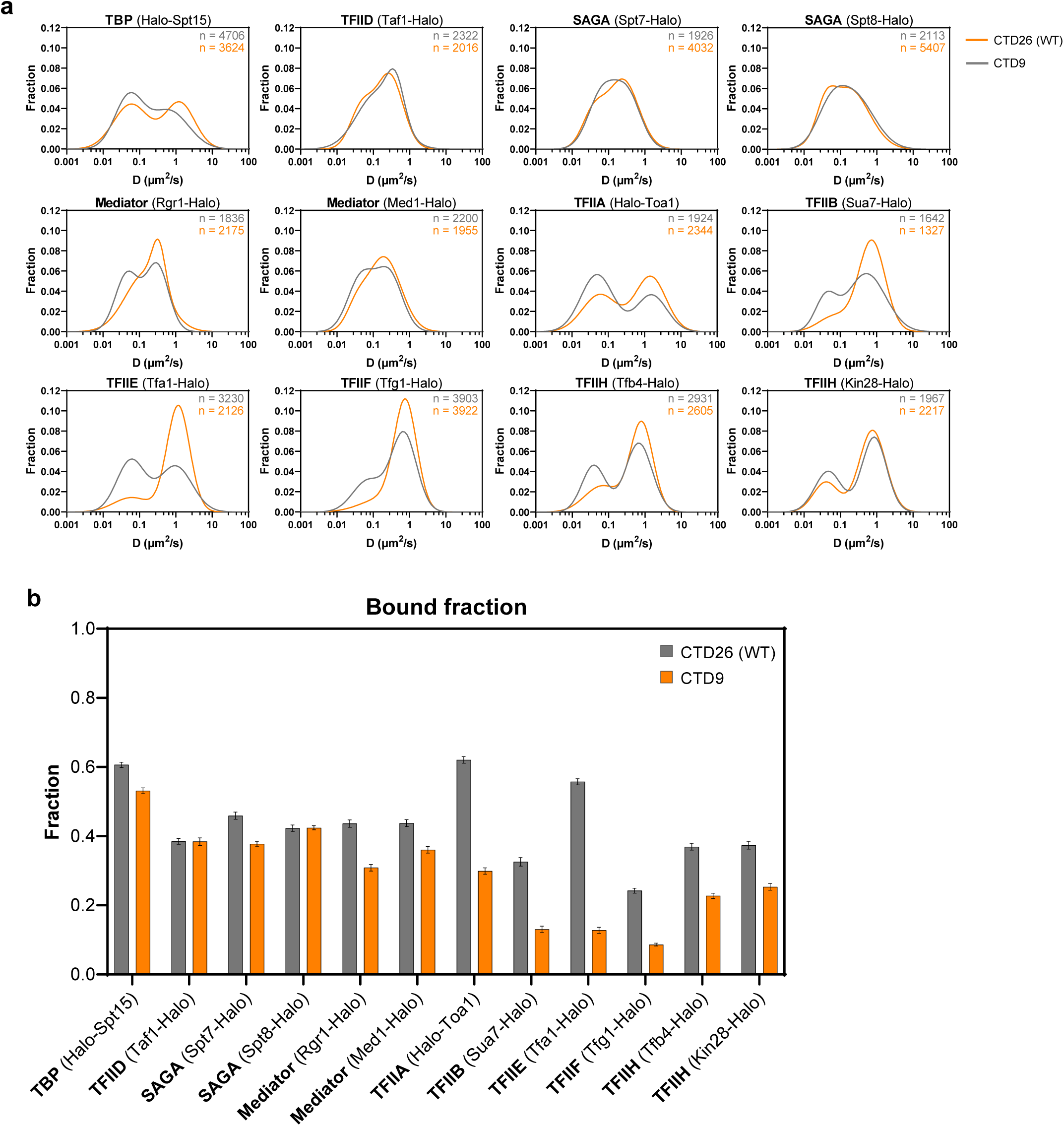
CTD truncation impairs PIC formation. **a**, Diffusion coefficient histogram for TBP (Halo-Spt15), TFIID (Taf1-Halo), SAGA (Spt7-Halo and Spt8-Halo), Mediator (Rgr1-Halo and Med1-Halo), TFIIA (Halo-Toa1), TFIIB (Sua7-Halo), TFIIE (Tfa1-Halo), TFIIF (Tfg1-Halo) and TFIIH (Tfb4-Halo and Kin28-Halo) in WT and CTD9 strains (n: number of trajectories). **b**, Fraction free and bound in WT and CTD9 strains (n = 100 resamplings; mean value ± s.d.).

We next examined the transcriptional regulation of heat shock response genes in the CTD9 mutant, due to its temperature-sensitive phenotype^7^. Hsf1, the master transcriptional regulator of the heat shock response genes^53^, constitutively associates with heat shock elements (HSEs) at the promoter^54–57^ under both non-heat shock (NHS) and heat shock (HS) conditions (Fig. 7f). The spatiotemporal map reveals a high density of bound Hsf1 near the nuclear envelope, similar to highly expressed Hsf1 gene targets in the genome simulation map (Fig. 7a,b). Likewise, the position of the highly expressed Hsf1 target gene *HSP104* exhibits a similar peripheral localization as determined by computational polymer modeling^58^. Hsf1 is highly dynamic, with over ∼85% of molecules that are freely diffusing (Fig. 7c and Extended Data Fig. 9a). Heat shock induction increases Hsf1 binding from ∼10% to ∼20% (stable binding from ∼2% to ∼5%) during the acute response (0–20 min at 39°C) (Fig. 7c,d), maintaining a ∼5 s residence time for NHS and acute HS conditions (Fig. 7e and Extended Data Fig. 9b). Prolonged HS (20–40 min at 39°C) slightly reduces Hsf1 binding fraction to ∼15% (∼3% stable binding) (Fig. 7c,d and Extended Data Fig. 9a), with a decrease of residence time to ∼2.5 s (Fig. 7e and Extended Data Fig. 9b). In the CTD9 mutant, Hsf1 binding and residence time during the acute heat shock response remains largely unchanged compared to the WT (Fig. 7c-e and Extended Data Fig. 9a,b), suggesting that CTD truncation has a minimal impact on the observed binding dynamics of the upstream transcription factor Hsf1.

**Fig. 7.**
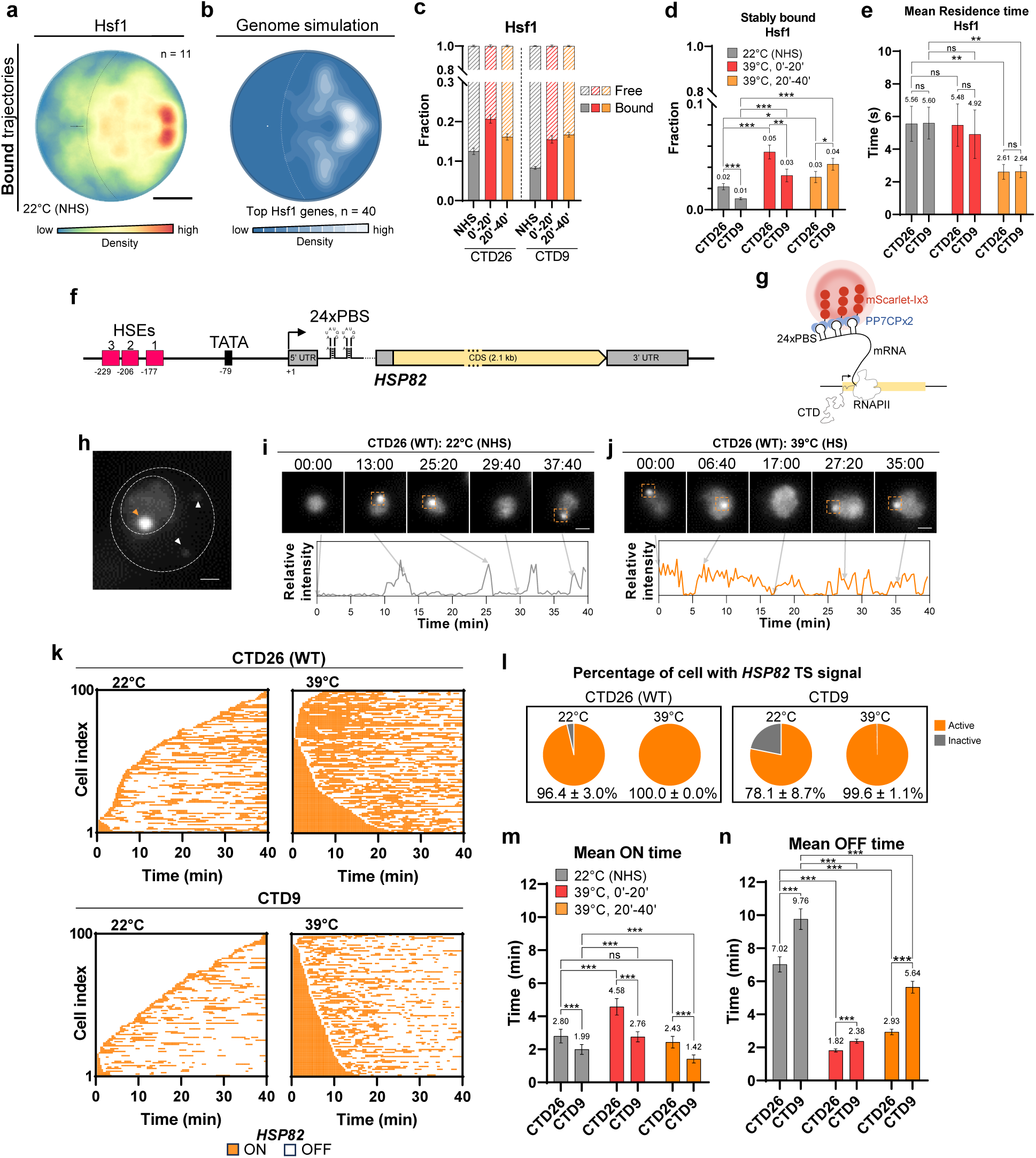
CTD length controls burst size and frequency of *HSP82* gene transcription. **a**, Spatiotemporal map of bound Hsf1-Halo trajectories. Scale bar: 0.5 µm. **b**, Simulation of top Hsf1 target genes (HSF1Base^70^; n: number of genes). **c**-**e**, **c**, Fraction free and bound (n = 100 resamplings; mean value ± s.d.), **d**, stably bound fractions (n = 10,000 resamplings; mean value ± s.d.; two-tailed unpaired t-test) and **e**, mean residence times (n = 10,000 resamplings; mean value ± s.d.; two-tailed unpaired t-test) of Hsf1-Halo on non-heat shock (HS) and heat shock (HS) for intervals of 0-20 minutes and 20-40 minutes. CTD9 mutant shows similar upregulation of Hsf1 binding during acute heat shock (39°C, 0’-20’) compared to WT, but with slightly lower basal levels. Unlike WT, the CTD9 mutant maintains Hsf1 binding at the induced level during the prolonged response (39°C, 20’-40’). The residence time of Hsf1 is comparable between WT and CTD9 under NHS and HS conditions. Overall, the dynamics of Hsf1 exhibit minimal change in CTD9 mutant. **f**, *HSP82* gene contains three characterized heat shock elements (HSEs) for Hsf1 binding^80^. 24x PP7 binding site (PBS) inserted within the 5’ UTR of *HSP82* gene. **g**, Nascent *HSP82* mRNA visualized by PP7CPx2-mScarlet-Ix3 binding. **h**, Nascent mRNAs on *HSP82* transcription site (TS) as a bright spot in the nucleus (orange arrow). Cytoplasmic *HSP82* mRNAs indicated by white arrows. Scale bar: 1.0 µm. **i**-**j**, Maximum z-projected time-lapse images of PP7-*HSP82* cells. Scale bar: 1.0 µm (top). Trace of relative fluorescence intensity of the transcription site (bottom). **k**, Binarized rastergrams of the PP7 signal at the *HSP82* TS of 100 cells, ordered by time of first detection. **l**, % cells with *HSP82* TS signal. **m**-**n**, **m**, Mean ON time and **n**, mean OFF time of *HSP82* transcription (n = 10,000 resamplings; mean value ± s.d.; two-tailed unpaired t-test).

To address the functional consequences of CTD truncation, we focused on *HSP82*, which is expressed constitutively under Hsf1 control and is highly inducible on heat shock. Previous research has emphasized the importance of CTD length in transcription activation^11–17^ and in the transcription bursting of *GAL* genes^13^, which are expressed only under inducible conditions. It is unclear whether the CTD controls the burst kinetics of constitutively expressed genes. We examined transcription bursting of *HSP82* by imaging the binding of fluorescently-tagged PP7 coat protein (PP7CPx2-mScarlet-Ix3) to a 24xPP7 sequence inserted into the 5’ UTR of the nascent *HSP82* mRNA (Fig. 7f,g and Supplementary Video 6). Strong fluorescent foci of PP7CPx2-mScarlet-Ix3 representing *HSP82* transcription sites (TSs), frequently appear near the nuclear envelope (Fig. 7h-j), consistent with the spatiotemporal map of bound Hsf1 (Fig. 7a,b). Under nonshock conditions, CTD9 shows a lower percentage of cells with *HSP82* TS fluorescence (∼78%) compared to WT (∼96%). However, upon heat shock, TS fluorescence appears almost instantaneously in nearly all cells for both WT and CTD9 mutant strains (Fig. 7k,l). We calculated the corresponding transcription ON and OFF times after binary conversion of *HSP82* TS signals (Fig. 7k). For WT CTD26, acute heat shock increases the *HSP82* ON time from ∼3 min to ∼4.5 min, while decreasing the OFF time from ∼7 min to ∼2 min. For prolonged heat shock, the ON time is similar to nonshock conditions, while the OFF time is reduced to ∼3 min (Fig. 7m,n, and Extended Data Fig. 9c,d), anticipating attenuation of the heat shock response. Notably, in all conditions, the CTD9 mutant consistently reduces the ON time (lower burst duration) and increases the OFF time (lower burst frequency) of *HSP82* transcription (Fig. 7m,n and Extended Data Fig. 9c,d), indicating that full CTD length is required for optimal burst kinetics under both constitutive and heat shock conditions. This finding offers a molecular explanation for the temperature-sensitive phenotype in the CTD9 mutant^7^. Importantly, the reduction in both duration and frequency of transcription bursting indicates that the diffusion confinement of RNAPII in the nucleoplasm could regulate the rate-limiting search for chromatin targets in the transcription cycle to sustain a transcription burst. The number of intrinsically disordered CTD repeats is central to this spatiotemporal confinement.

## Discussion

We have extended single-molecule tracking (SMT) to develop global spatiotemporal maps of transcription protein dynamics in living yeast (Fig. 1). This approach reveals that the disordered CTD of RNAPII, with a WT length of 26 heptad repeats, directs the confinement of RNAPII within a subnuclear region enriched for active genes (Fig. 2). Characterization of anisotropic diffusion of RNAPII shows its dependence on the full-length CTD, and on chromatin organization and the presence of Mediator (Fig. 3). RNAPII exhibits transient confinement without phase separation into condensates (Fig. 4). Importantly, the number of CTD repeats critically influences RNAPII search kinetics (Fig. 5) and PIC formation (Fig. 6). Furthermore, CTD truncation functionally disrupts transcription burst kinetics of *HSP82* (Fig. 7).

Our observations in yeast demonstrate that RNAPII does not exhibit substantial clustering, questioning the generality and the extent of LLPS involvement in its spatial organization. For eukaryotes with larger nuclei and genome complexity, additional spatial control and dynamic phase-separated compartments might be necessary for long range promoter-enhancer interaction^59,60^, although this is a topic of ongoing study. In yeast, however, the strong physical constraints and subnuclear chromosome tethering^32–35^, alongside the compact genome, could create domains enriched for active genes (Fig. 1c,d), and facilitate RNAPII transient confinement without the need for phase separation into condensates (Fig. 3 and 4). Furthermore, the concentration of RNAPII in yeast (1,200 molecules per µm^3^, assuming a nuclear radius of 1 µm, and 5,000 RNAPII per nucleus (least abundance subunit Rpb9; SGD)) is ∼30-fold higher than in human U2OS cell nuclei (∼40 molecule per µm^3^)^61^. Such a high endogenous RNAPII concentration in yeast suggests that transient confinement might be sufficient for efficient target search. Furthermore, the yeast nucleoplasmic environment may be less favorable for phase separation into condensates compared to higher eukaryotes.

What might be the molecular basis of the spatiotemporal confinement of yeast RNAPII? Weak interactions observed *in vitro* between the IDRs of the RNAPII CTD and those of the Mediator complex^20–22^ (Supplementary Fig. 3), hint at a potential mechanism involving IDR interactions that remains to be further elucidated. Imaging of the CTD fused to GFP shows evidence of enrichment at transcriptionally active loci of Drosophila polytene chromosomes *in vivo*^62^, raising the question whether the CTD alone is sufficient to drive its own spatiotemporal confinement in living yeast. Our supplementary observations show that HaloTag alone and Halo-CTD fusions harboring 26, 52, 78, and 104 heptad repeats expressed under control of the native *RPO21* promoter exhibit minimal chromatin binding (Extended Data Fig. 10a), show no puncta formation (Extended Data Fig. 10b), and D_free_ values aligning with the Stokes-Einstein equation, indicating uniform nucleoplasmic viscosity (∼30 cP) without phase separation (Extended Data Fig. 10d-g). Furthermore, Halo-CTD mutants do not exhibit high f_180/0_ values, indicating that diffusion is relatively isotropic or unconfined in the yeast nucleus (Extended Data Fig. 10h). Evidently, at physiological expression levels, the CTD alone does not induce the same degree of diffusion confinement as observed for the RNAPII complex. Thus, the CTD is necessary, but not sufficient for the spatiotemporal confinement of RNAPII, and additional factors or domains within the RNAPII complex might come into play.

In addition to a structural role of the CTD in driving PIC formation^49,50,52^, our data suggest that transient spatiotemporal confinement of freely diffusing RNAPII dependent on CTD repeats and the Mediator facilitates the rate-limiting search for chromatin targets early in the transcription cycle (Fig. 8). In particular, diffusion confinement of RNAPII could increase the probabilities of intermolecular interactions and specific association rates with the Mediator and other PIC components, promoting repeated rounds of transcription to generate a transcription burst. Our study emphasizes the crucial role of the CTD in driving RNAPII target search and shaping the spatiotemporal dynamics of PIC assembly, highlighting its multifaceted functions in the pathway to gene expression.

**Fig. 8.**
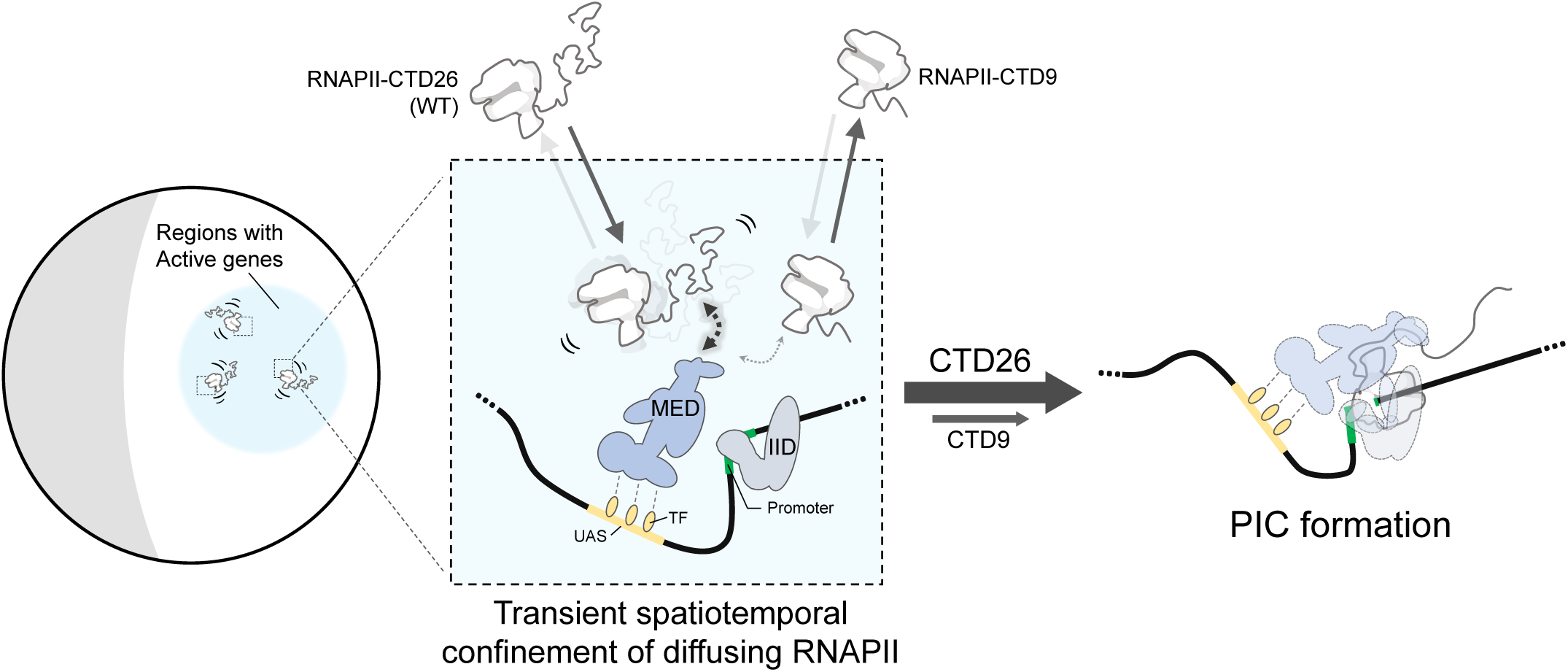
CTD directs subnuclear confinement and chromatin binding of RNAPII for efficient PIC assembly and transcription bursting. In the yeast nucleus, physical constraints such as telomere tethering of yeast chromosomes create subnuclear regions enriched with active genes (left blue circle). Within these regions, the diffusion of RNAPII is transiently confined, potentially through weak and long-range interaction (dashed arrows) between the CTD and the Mediator near an active promoter (middle square), leading to localized oversampling and enhanced target search efficiency. Such subnuclear confinement of freely diffusing RNAPII promotes the probability of intermolecular interactions and specific association rates for the formation of the multi-component PIC, and facilitates multiple rounds of RNAPII binding and activation for a transcription burst. IID: TFIID; MED: Mediator; TF: Transcription factor; UAS: Upstream activating sequence.

## Acknowledgments

We thank Scarlet Cho and Gan Ling for assistance in image processing; Luke Lavis for providing Janelia Fluor dyes; Kelly Xie for IUPRED3 data mining; Xiaona Tang and Sheng Liu for computational assistance; Nathan Jones, Vincent Schoonderwoert and Ciprian Almonte for microscopy technical support; Karen Yuen and Gaku Mizuguchi for plasmids and yeast strains; Craig Kaplan and Joanna Yao for valuable information on strains sensitivity on MPA; Anders Hansen, Andrea Musacchio, Charmaine Wong, Ibrahim Cissé and Jee Min Kim for helpful discussions; and members of the Wu laboratory for comments. This study was supported by National Institute of Health grant GM132290-01 (C.W.) and the Croucher Foundation (Y.H.L.).

## Author information

These authors contributed equally: Ziyang Ye, Chloe Liang, Chuofan Yu, Giho Park

## Contributions

Y.H.L., J.L.C., and C.W. designed the experiments. Y.H.L. performed the experiments and analyzed the data. Z.Y. performed SMT on Hsf1-Halo in the wildtype background. G.P. contributed to the initial trial of spatiotemporal mapping. Z.Y., C.L., and C.Y. assisted in experiments. Y.H.L., J.L.C., and C.W. wrote the manuscript.

## Methods

### Extended Data Figures

Extended Data Figures 1-10

### Supplementary information

Supplementary Methods, Supplementary Tables 1,2, Supplementary Figures 1-4 and references. Supplementary Videos 1-6

---

**Extended Data Fig. 1.**
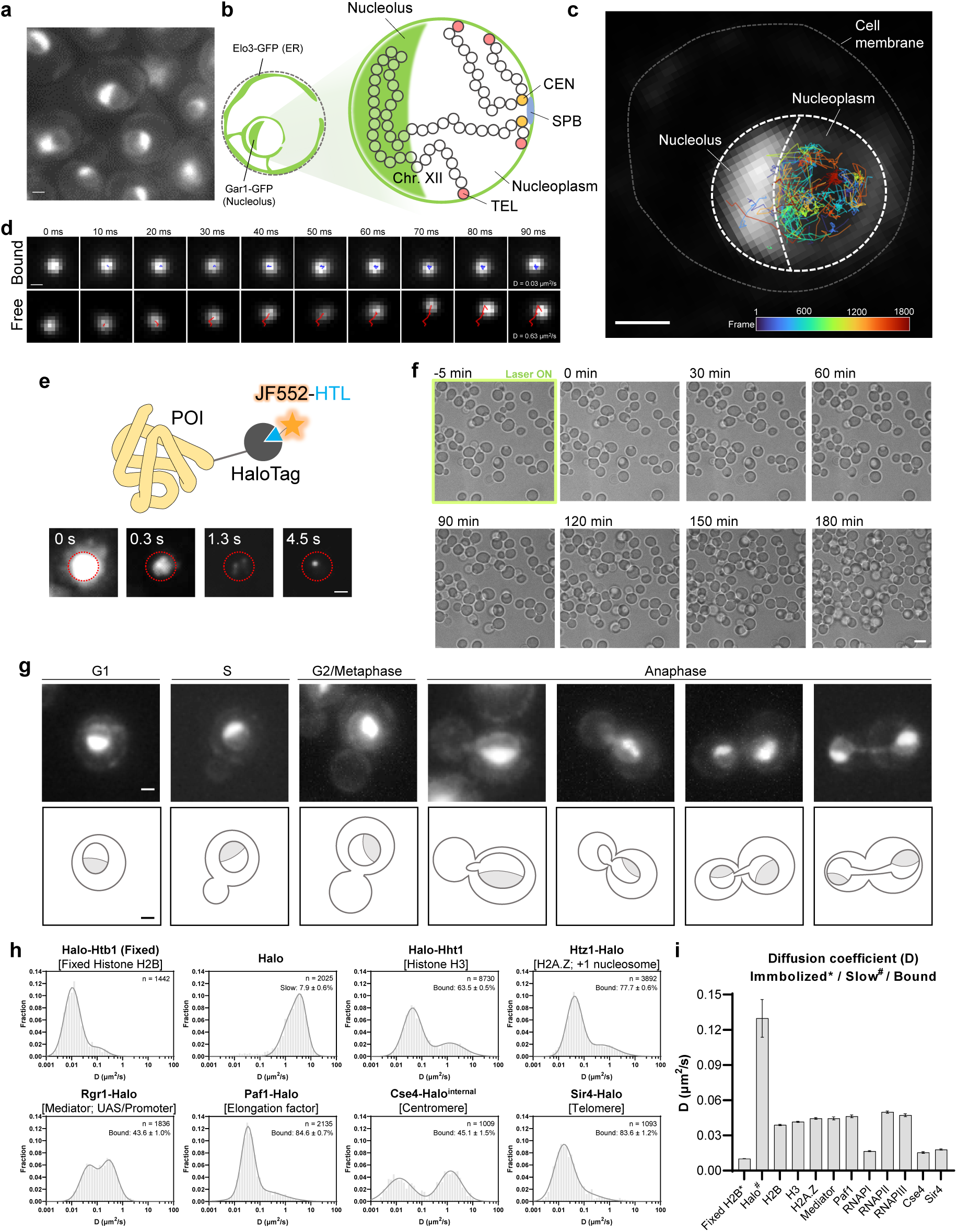
Single-molecule tracking in living yeast. **a**, ER and nucleolus shown by Elo3-GFP and Gar1-GFP. Scale bar: 1.0 µm. **b**, Coarse-grained representation of yeast interphase chromosomes, physically constrained by telomere (TEL)-nuclear envelope and centromere-spindle pole body (CEN-SPB) tethering. **c**, Overlay of RPB1^Rpo21^-Halo^JF552^ trajectories on the nucleus relative to ER and nucleolar GFP markers. Rainbow colors indicate the first appearance of each trajectory. Scale bar: 1 µm. **d**, Single-molecule trajectories of RPB1^Rpo21^-Halo^JF552^ (free or bound) with diffusion coefficients. Scale bar: 0.2 µm. **e**, Protein of interested (POI) fused with HaloTag, labeled with JF552-HaloTag ligand (JF552-HTL) (top). Initial laser exposure of JF552 resulted in strong nuclear glow’, followed by shelving in dark state and stochastic reactivation, leading to single-molecule detection. Nucleus circled in red. Scale bar: 1.0 µm. (bottom). **f**, Strong 555 nm laser exposure for 5 min does not result in noticeable cell death or growth arrest. **g**, ER and nucleolar GFP markers for cell cycle stage identification. Scale bar: 1.0 µm. **h**, Diffusion coefficient histograms of various nuclear proteins (n: number of trajectories; mean value ± s.d.). **i**, Assessing diffusion coefficients of the immobilized/slow/bound population reveals no significant difference between histones and proteins that can move along DNA (e.g. RNAPII, RNAPIII, Paf1), while nucleolar RNAPI, and proteins that are physically tethered (Cse4 and Sir4) have significantly lower values (n = 100 resamplings; mean value ± s.d.).

**Extended Data Fig. 2.**
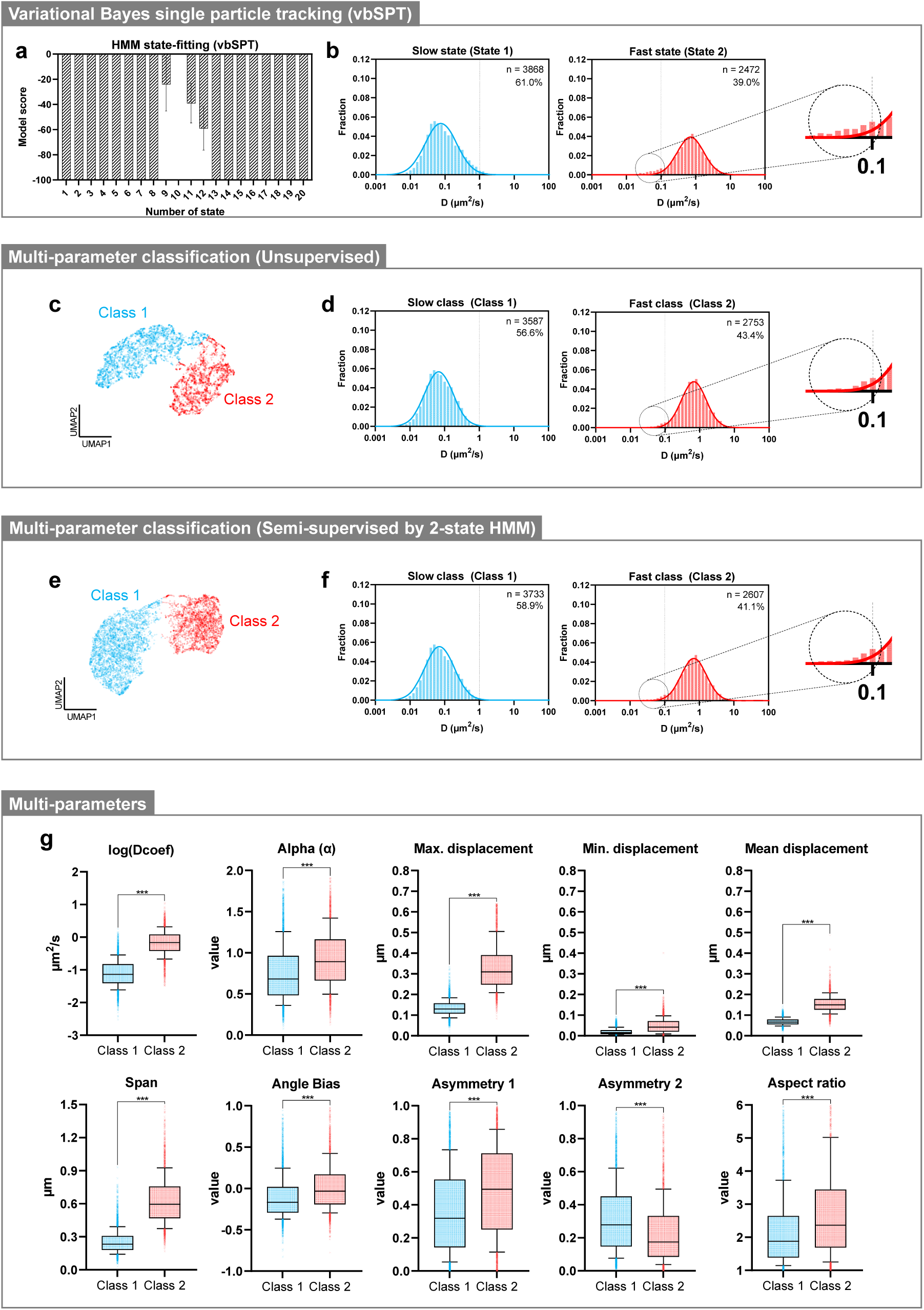
Multi-parameter classification of trajectories. **a**, Unsupervised vbSPT overfits SMT data of RNAPII (n = 100 resamplings; mean value ± s.d.). **b**, Diffusion coefficient histogram for fast and slow states from 2-state HMM using vbSPT. A minor fraction of very slowly diffusing trajectories was misclassified as fast-state (dashed circle) (n: number of trajectories; mean value ± s.d.). **c**, Unsupervised multi-parameter classification of RNAPII trajectories with UMAP dimensionality reduction and GMM clustering. **d**, Diffusion coefficient (D) distribution for the classified clusters (n: number of trajectories; mean value ± s.d.). **e**-**f**, similar to **c**-**d**, but with multi-parameter classification semi-supervised by 2-state HMM. **d** and **f**, Compared to **b**, Fast-class diffusion coefficient distribution showed less tailing (dashed circle). **g**, 10 parameters used for classification and their distributions in identified classes (refers **f** for n numbers; median (line), IQR (box), 10-90 percentile (whiskers) and outliers (crosses); two-tailed unpaired t-test).

**Extended Data Fig. 3.**
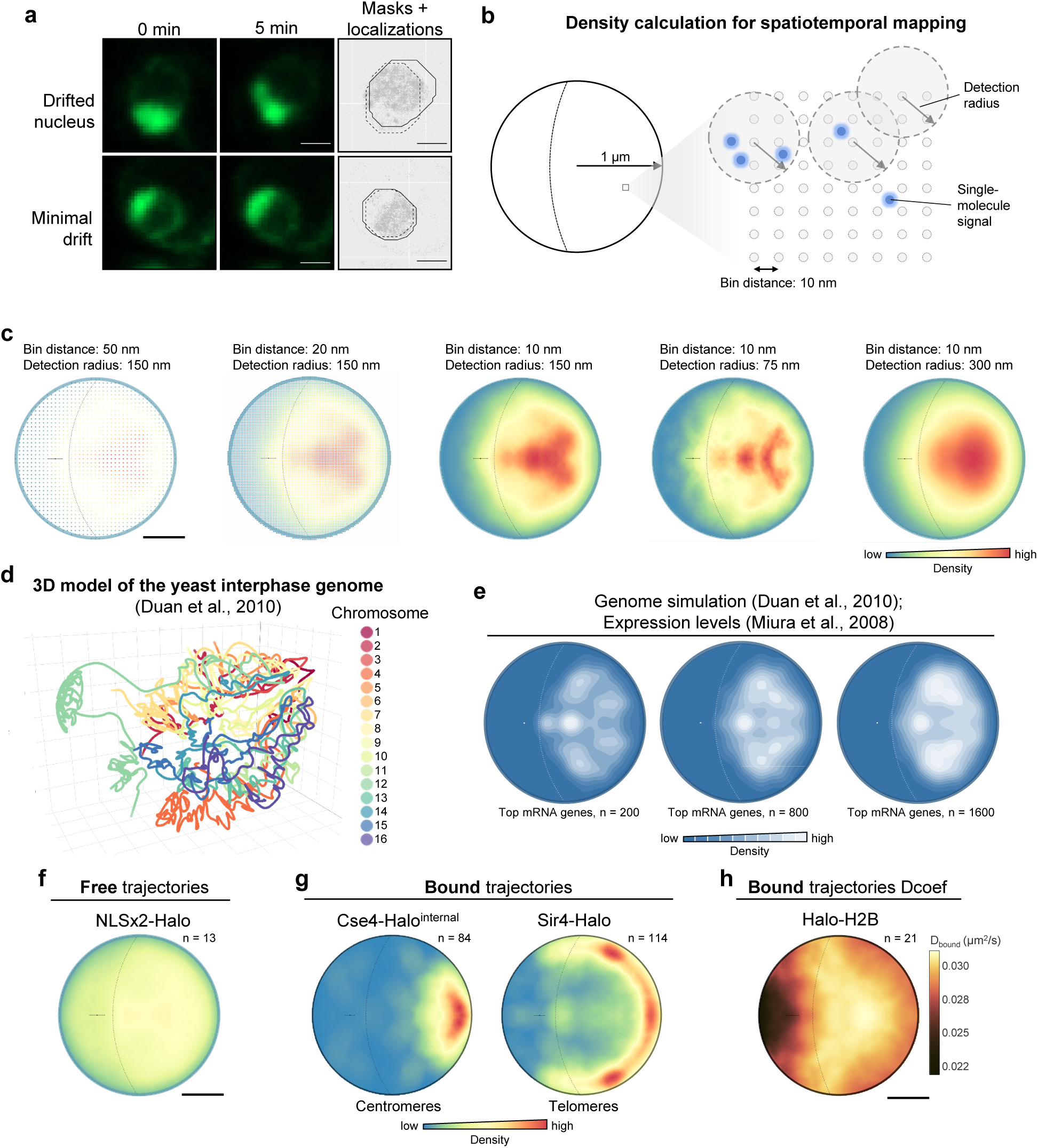
Spatiotemporal mapping reveals dynamics of nucleosome and nuclear landmark proteins. **a**, Nuclei in G1 cells with minimal drift selected for spatiotemporal mapping. Scale bar: 1.0 µm. **b**, Localization density calculated using 150 nm detection radius from bins separated by 10 nm. **c**, 150 nm detection radius and bin distance of 10 nm provide optimal coverage and resolution for the density on the bound map of RNAPII. Scale bar: 0.5 µm (Error bar: centroid of nucleolus ± s.d.). **d**, 3D simulation of the yeast interphase genome based on 3C data^37^. **e**, Top mRNA genes simulation maps based on gene expression data in minimal medium^38^. Left panel is reuse of Fig. 1d top panel (n: number of genes). **f**, Freely diffusing NLSx2-Halo map. Scale bar: 0.5 µm. **g**, Bound Cse4 (left) and Sir4 (right) maps (Error bar: centroid of nucleolus ± s.d.; n: number of nuclei). Sir4 bound map showed dense regions proximal and distal to the centromere, mirroring the telomere clusters of short and long chromosomes^35,37,81^. **h**, Bound H2B diffusion coefficient map. A 300 nm detection radius was used. Scale bar: 0.5 µm (Error bar: centroid of nucleolus ± s.d.; n: number of nuclei). Telomere and centromere movements are confined due to attachment to the nuclear envelope and spindle pole body, respectively^82^. Likewise, the local diffusion coefficients of chromatin-bound H2B reveal lower histone dynamics near centromeres and telomeres (Extended Data Fig. 1h,i).

**Extended Data Fig. 4.**
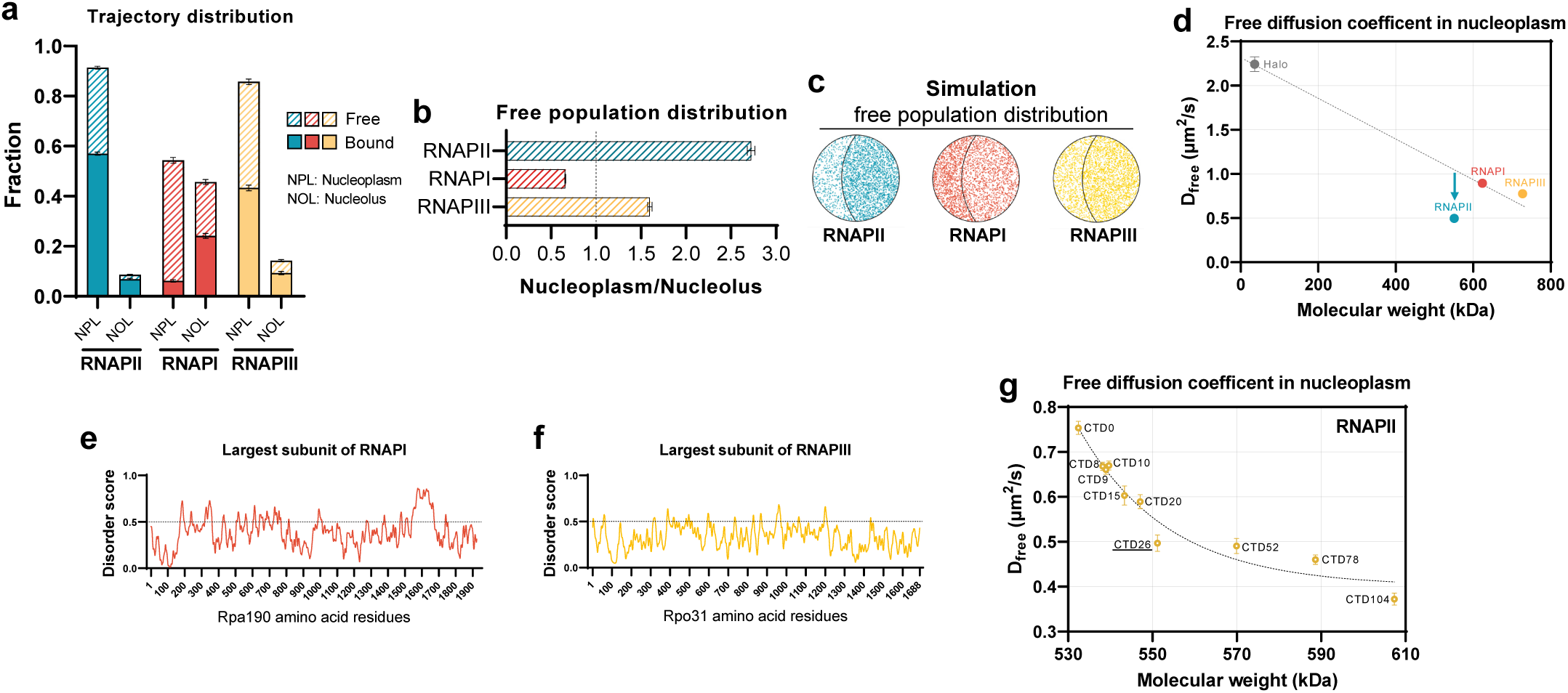
Spatiotemporal dynamics of RNAPII, RNAPI and RNAPIII. **a**, Fraction of free and bound in nucleoplasm (NPL) and nucleolus (NOL) (n = 100 resamplings; mean value ± s.d.). **b**, Distribution (area normalized) of free RNAPII, RNAPI and RNAPIII in nucleoplasm and nucleolus (n = 100 resamplings; mean value ± s.d.). **c**, Simulation of Nucleoplasm/Nucleolus distribution. **d**, Diffusion coefficients and molecular weights of free RNAPII, RNAPI and RNAPIII in nucleoplasm (n = 100 resamplings; mean value ± s.d.). **e**-**f** IUPRED3 disorder scores for the largest subunit of **e**, RNAPI (Rpa190) and **f**, RNAPIII (Rpo31). Residues with predicted scores above 0.5 considered disordered. **g**, D_free_ of RNAPII and CTD mutants in nucleoplasm (n = 100 resamplings; mean value ± s.d.).

**Extended Data Fig. 5.**
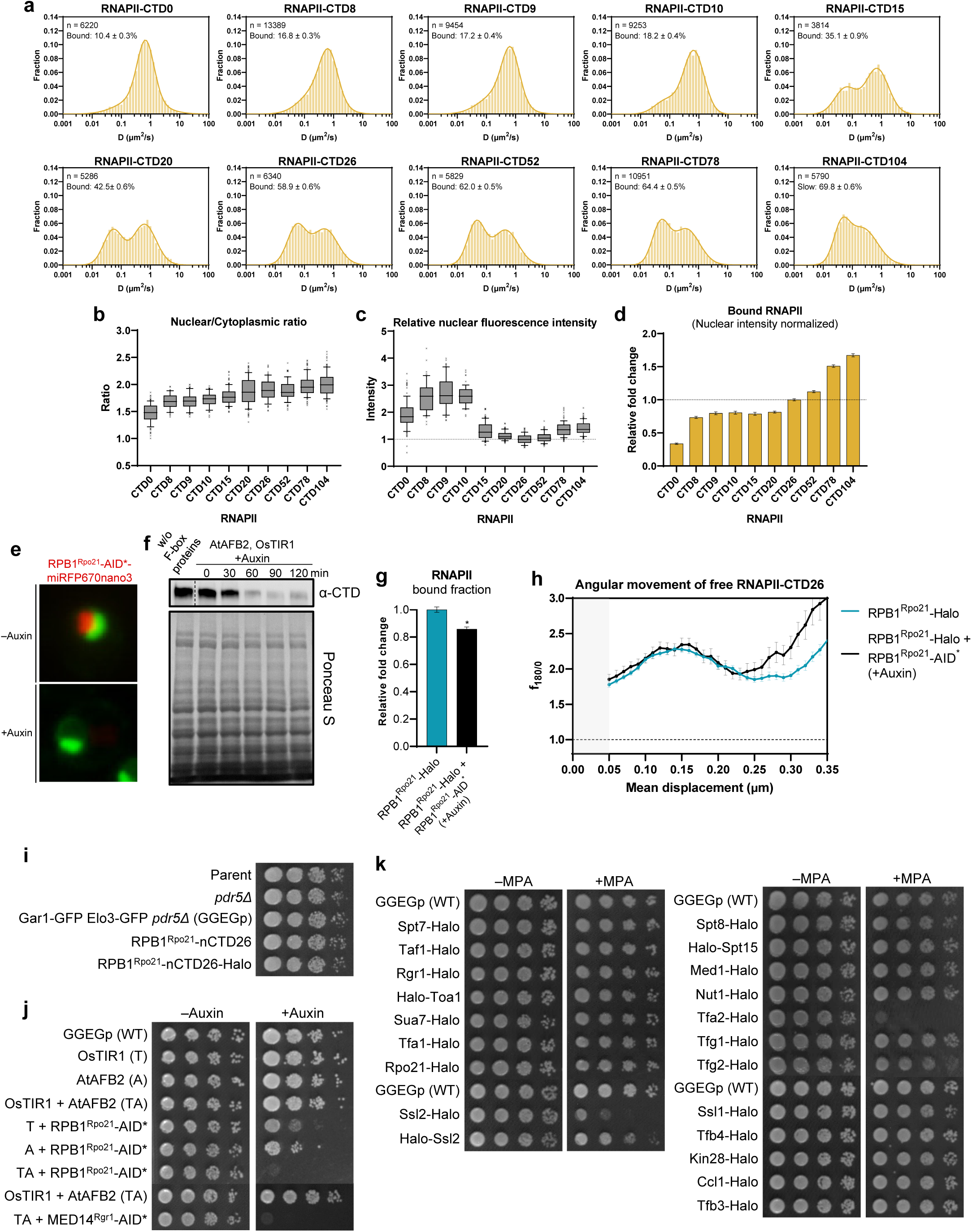
CTD length affects RNAPII subcellular distribution and expression. **a**, Diffusion coefficient histogram for RNAPII with varying CTD lengths (n: number of trajectories; mean value ± s.d.). **b**-**c**, **b**, Nuclear/Cytoplasmic ratio and **c**, relative nuclear intensity of WT RNAPII and CTD mutants (n ≥ 50 nuclei; median (line), IQR (box), 10-90 percentile (whiskers) and outliers (crosses)). CTD truncation led to increased cytoplasmic and nucleolar distribution (Fig. 2b), and overexpression of RNAPII. **d**, Amount of bound RNAPII normalized to nuclear fluorescence intensity (n = 100 resamplings; mean value ± s.d.). The normalized value is an approximation, as the relationship between nuclear fluorescence intensity and RNAPII levels may not be strictly linear. **e**, Fluorescence signal of RPB1^Rpo21^-AID*-miRFP670nano3 (red) before and after RNAPII degradation by auxin. Nucleolar and ER markers in green. Scale bar: 1.0 µm. **f**, Western blot showing time course degradation of RPB1^Rpo21^-AID* by auxin, detected using CTD antibody (top), and Ponceau S staining on total protein (bottom). **g**-**h**, Cells expressing ectopic RPB1^Rpo21^-Halo in an RPB1^Rpo21^ AID background show minimal changes in the **g**, bound fraction (n = 100 resamplings; mean value ± s.d.; two-tailed unpaired t-test) and **h**, f_180/0_ of RNAPII after auxin treatment, compared to those in WT expressing RPB1^Rpo21^-Halo. **i**, Spot assay for strains with *pdr5*Δ, GFP markers (GGEGp), RPB1^Rpo21^-nCDT26, or RPB1^Rpo21^-nCDT26-Halo showing identical growth. **j**, Spot assay demonstrating impaired cell growth for RPB1^Rpo21^ or MED14^Rgr1^ AID degron. All AID* constructs contain miRFP670nano3 as a fluorescence marker. **k**, Spot assay for strains with HaloTag fusions to various PIC component subunits to assess MPA sensitivity. Ssl2-Halo (TFIIH), Halo-Ssl2 (TFIIH), Tfa2-Halo (TFIIE), Tfg2-Halo (TFIIF) exhibited MPA sensitivity. Rpo21-Halo in **k** is identical to RPB1^Rpo21^-nCTD26-Halo in **i**.

**Extended Data Fig. 6.**
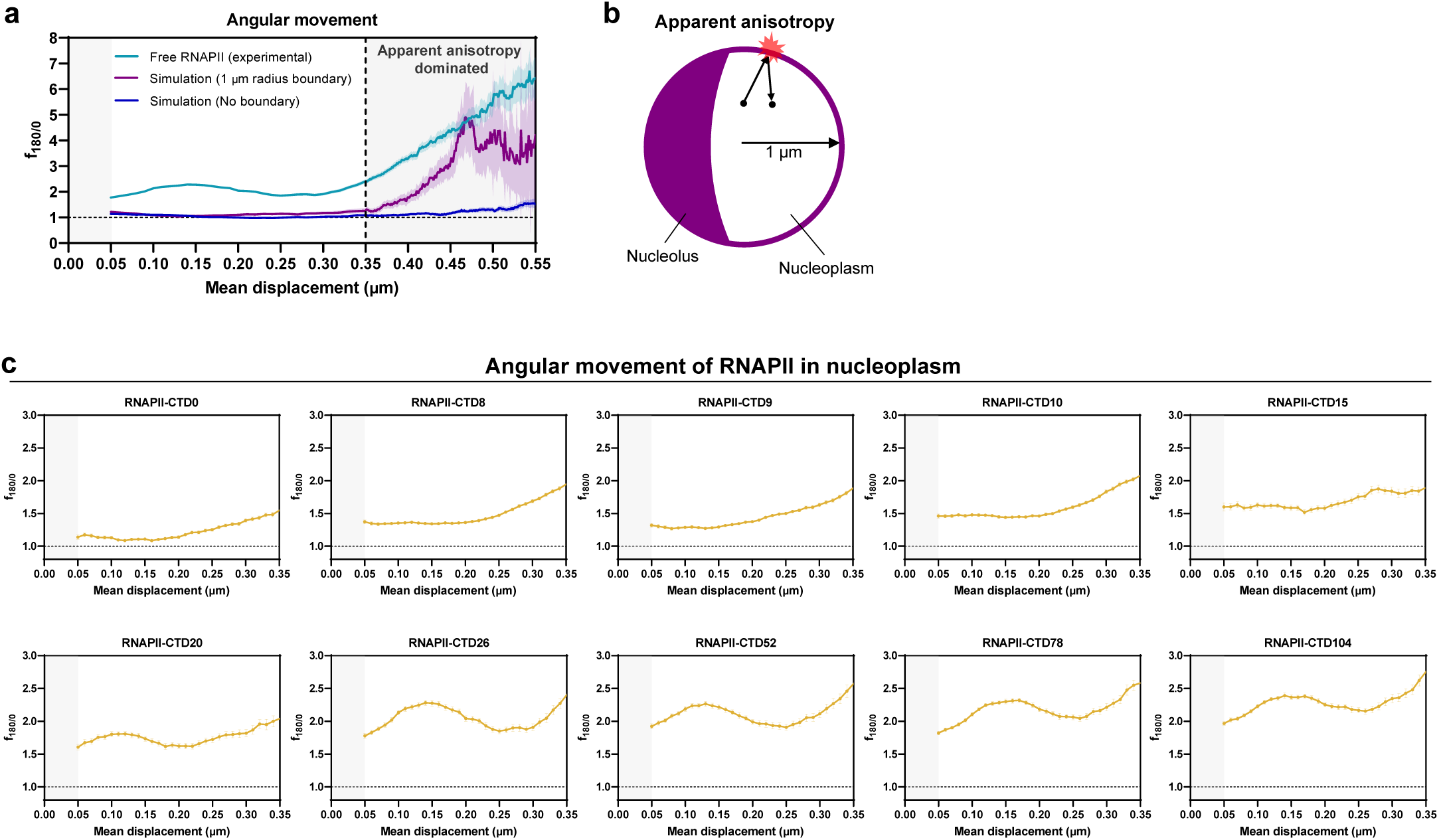
CTD length impacts free RNAPII anisotropy. **a**, Brownian simulations indicate significant apparent anisotropy for mean displacement longer than 350 nm (n = 100 resamplings; mean value ± s.d.). **b**, Apparent anisotropy due to nuclear confinement in small yeast nucleus. **c**, CTD length increase leads to an elevation in f_180/0_ trend (n = 100 resamplings; mean valu ± s.d.).

**Extended Data Fig. 7.**
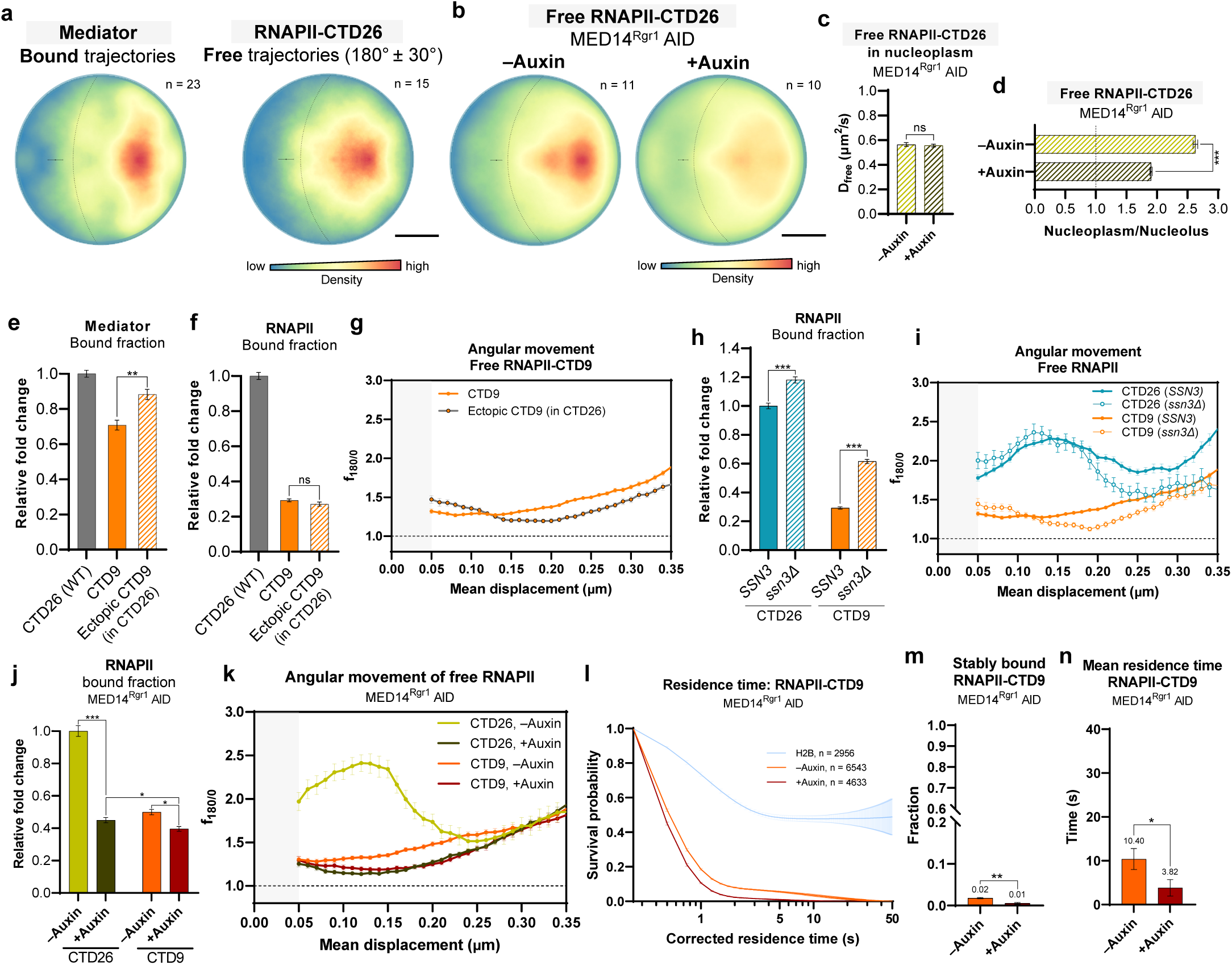
Investigating the role of Mediator in spatiotemporal confinement of RNAPII. **a**, Bound Mediator map (left). Free RNAPII-CTD26 map showing confined trajectories (right). Scale bar: 0.5 µm. (Error bar: centroid of nucleolus ± s.d.; n: number of nuclei). **b**, Free RNAPII-CTD26 map in MED14^Rgr1^ AID background before and after auxin treatment. Scale bar: 0.5 µm (Error bar: centroid of nucleolus ± s.d.; n: number of nuclei). **c**-**d**, **c**, D_free_ in nucleoplasm and **d**, Nucleoplasm/Nucleolus value of RNPAII-CTD26 in MED14^Rgr1^ AID background before and after auxin treatment (n = 100 resamplings; mean value ± s.d.; two-tailed unpaired t-test). The large hydrodynamic volume due to the disordered CTD of RNAPII (Extended Data Fig. 10d,e) may hinder its entry into the nucleolus by size exclusion (Fig. 2c, Extended Data Fig. 4b, and 10c). Additionally, Mediator degradation increases the diffusion of WT RNAPII-CTD26 into the nucleolus. This underscores the role of Mediator in maintaining RNAPII confinement within the nucleoplasmic space. **e**-**g**, **e**, Relative Mediator bound fraction, **f**, relative RNAPII bound fraction and **g**, f_180/0_ for RNAPII-CTD9 as sole source, or ectopically expressed in CTD26 (WT) background (n = 100 resamplings; mean value ± s.d.; two-tailed unpaired t-test). A moderate decrease in Mediator binding (Rgr1-Halo and Med1-Halo) observed in the CTD9 mutant (Fig. 6a,b) prompted us to examine if this could account for the observed RNAPII-CTD9 dynamics (Fig. 2 and 3). Despite the restoration of Mediator binding (Rgr1-Halo) in a strain co-expressing RPB1^Rpo21^-CTD26 (WT) and RPB1^Rpo21^-CTD9, RNAPII-CTD9 consistently exhibited a significant decrease in bound fraction and near-isotropic diffusion. This suggests that the changes in RNAPII dynamics are more likely intrinsic to the CTD truncation, rather than being predominantly driven by the decreased Mediator binding. **h**, Relative bound fraction of RNAPII-CTD26 and RNAPII-CTD9 in WT and *ssn3*Δ (n = 100 resamplings; mean value ± s.d.; two-tailed unpaired t-test). **i**, f_180/0_ plot for RNAPII-CTD26 and RNAPII-CTD9 in WT and *ssn3*Δ (n = 100 resamplings; mean value ± s.d.). **h**-**i**, During PIC formation, interaction between the Mediator and RNAPII can be obstructed by the Mediator regulatory subcomplex, the 4-subunit Cdk8 kinase module (CKM)^83–85^. Deficiency of Cdk8*^SSN3^* (*ssn3*Δ) promotes stable Mediator-RNAPII interactions^83^. Importantly, increasing the frequency of stable interactions between Mediator and RNAPII does not affect the confinement of free RNAPII, as little change in the anisotropy (f_180/0_) of free RNAPII is observed in the *ssn3*Δ mutant. **j**-**k**, **j**, Relative bound fraction and **k**, f_180/0_ plot for RNAPII-CTD26 and RNAPII-CTD9 in MED14^Rgr1^ AID background before and after auxin treatment (n = 100 resamplings; mean value ± s.d.; two-tailed unpaired t-test). **l**-**n**, **l**, Survival probability of H2B-corrected residence times, **m**, stably bound fractions and **n**, mean residence times for RNAPII-CTD9 in MED14^Rgr1^ AID background before and after auxin treatment (n = 10,000 resamplings; mean value ± s.d.; two-tailed unpaired t-test).

**Extended Data Fig. 8.**
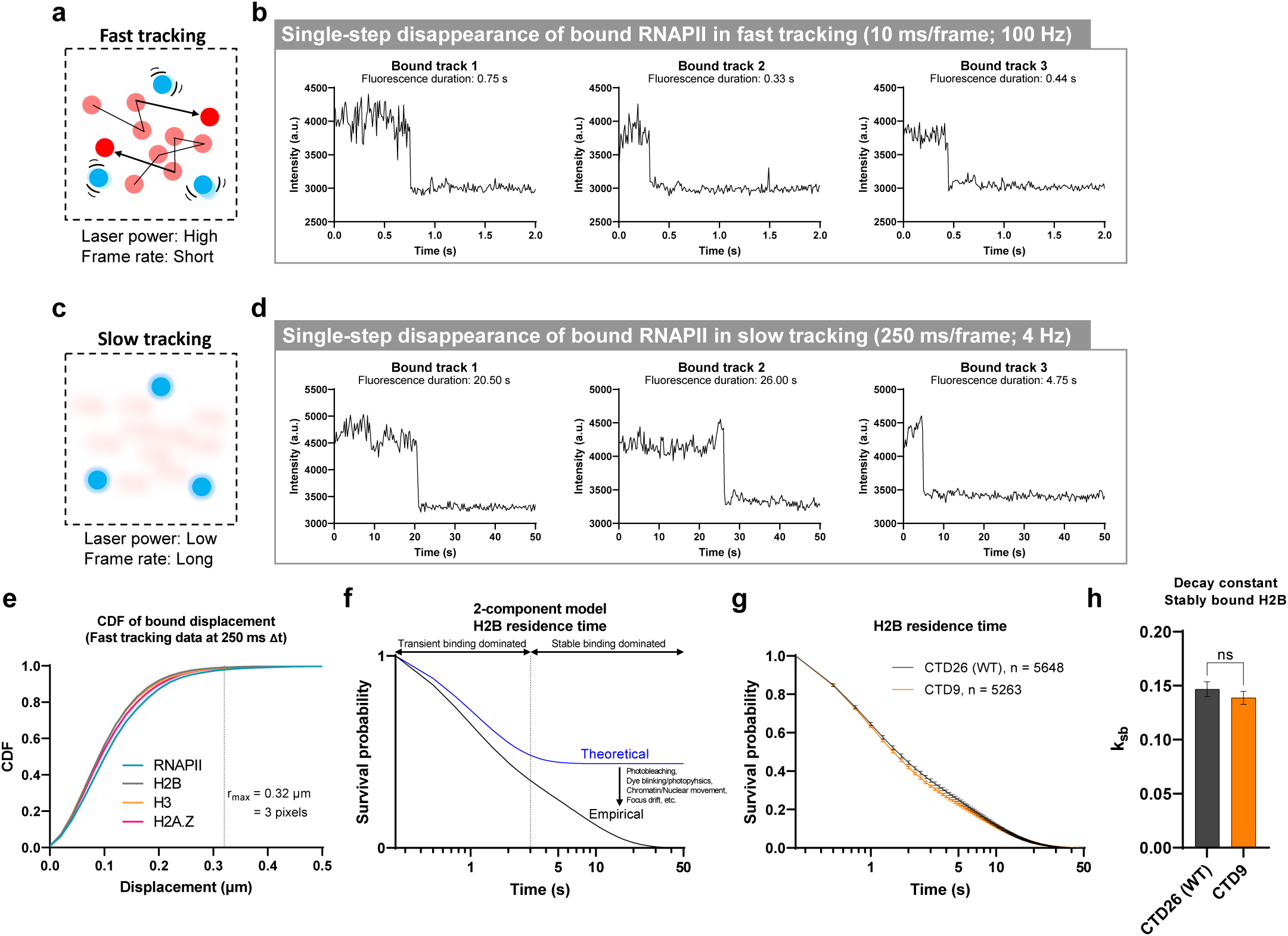
Fast and slow tracking mode of single-molecule tracking. **a**, Fast tracking with high laser power and short frame rate (10 ms/frame) reveals diffusive dynamics of free (red) and bound (blue) molecules. **b**, Single-step disappearance of chromatin-bound RNAPII in fast tracking. **c**, Slow tracking with low laser power and long frame rate (250 ms/frame) reveals residence time of bound molecules (blue). **d**, Single-step disappearance of chromatin-bound RNAPII in slow tracking. **b**-**d**, Here we term disappearance’ as we cannot definitively differentiate between the molecule diffusing out of the focal plane or photobleaching within the living cell. **e**, CDF of bound displacement of RNAPII, H2B, H3 and H2A.Z to determine the r_max_ for trajectory linking in slow tracking^86^. **f**, In G1 cells, nucleosomal H2B (representing the stable binding population) was hypothesized to bind for a sufficiently long period, such that during image acquisition, their dissociation was barely observed. Nonetheless, the observed residence times for H2B typically fall within 30 seconds, due to factors such as photobleaching, dye blinking, chromatin/nuclear movements, and focus drift. **g**, Survival probability of Halo-H2B observed residence times in CTD26 (WT) and CTD9 strains (n: number of trajectories; mean value ± s.d.). **h**, Decay constant of stably bound Halo-H2B (k_sb_) in CTD26 (WT) and CTD9 strains (n = 10,000 resamplings; mean value ± s.d.; two-tailed unpaired t-test).

**Extended Data Fig.9.**
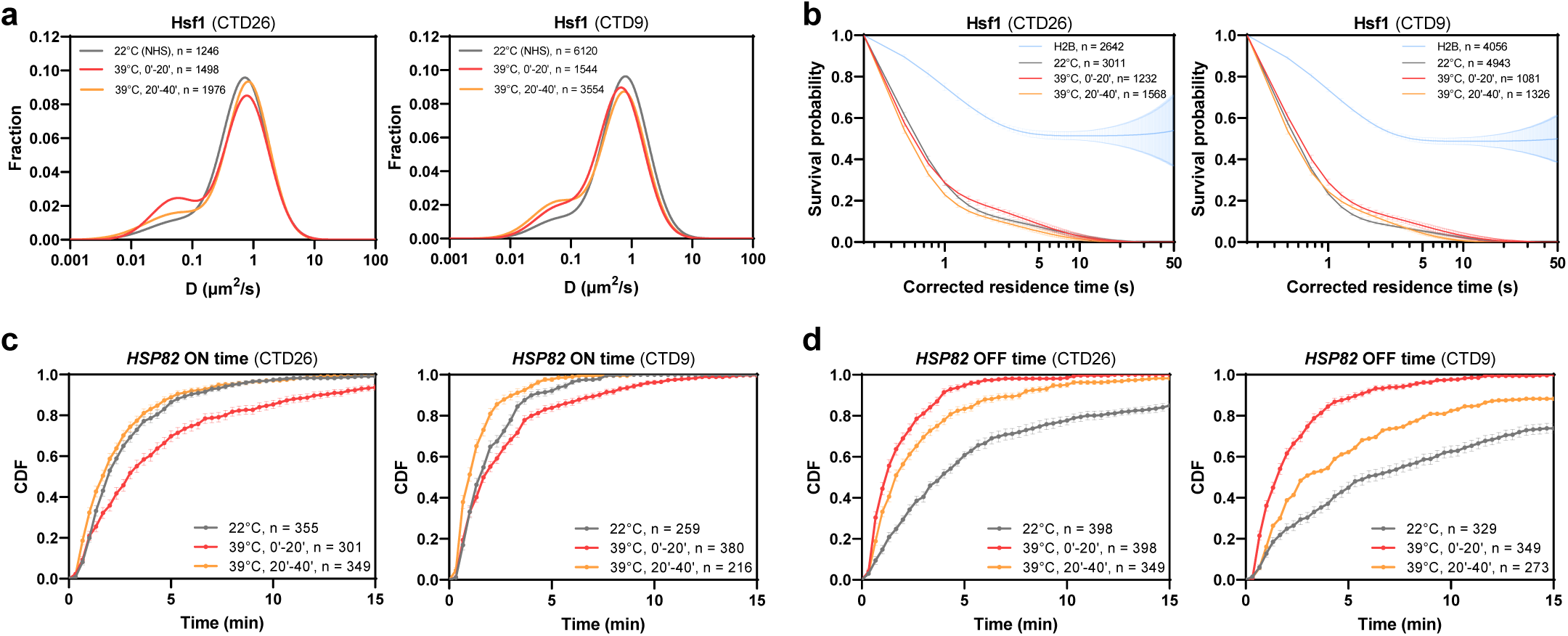
CTD length controls *HSP82* transcription bursting, with minimal effect on Hsf1 dynamics. **a**-**b**, **a**, Diffusion coefficient histograms for Hsf1-Halo and **b**, survival probability of bound Hsf1-Halo under heat shock in CTD26 (WT) (left) and CTD9 (right) strains (n: number of trajectories; mean value ± s.d.). **c**-**d**, **c**, *HSP82* ON time and **d**, OFF time cumulative distribution function (CDF) under heat shock in CTD26 (WT) (left) and CTD9 (right) strains (n: number of ON or OFF events; mean value ± s.d.).

**Extended Data Fig. 10.**
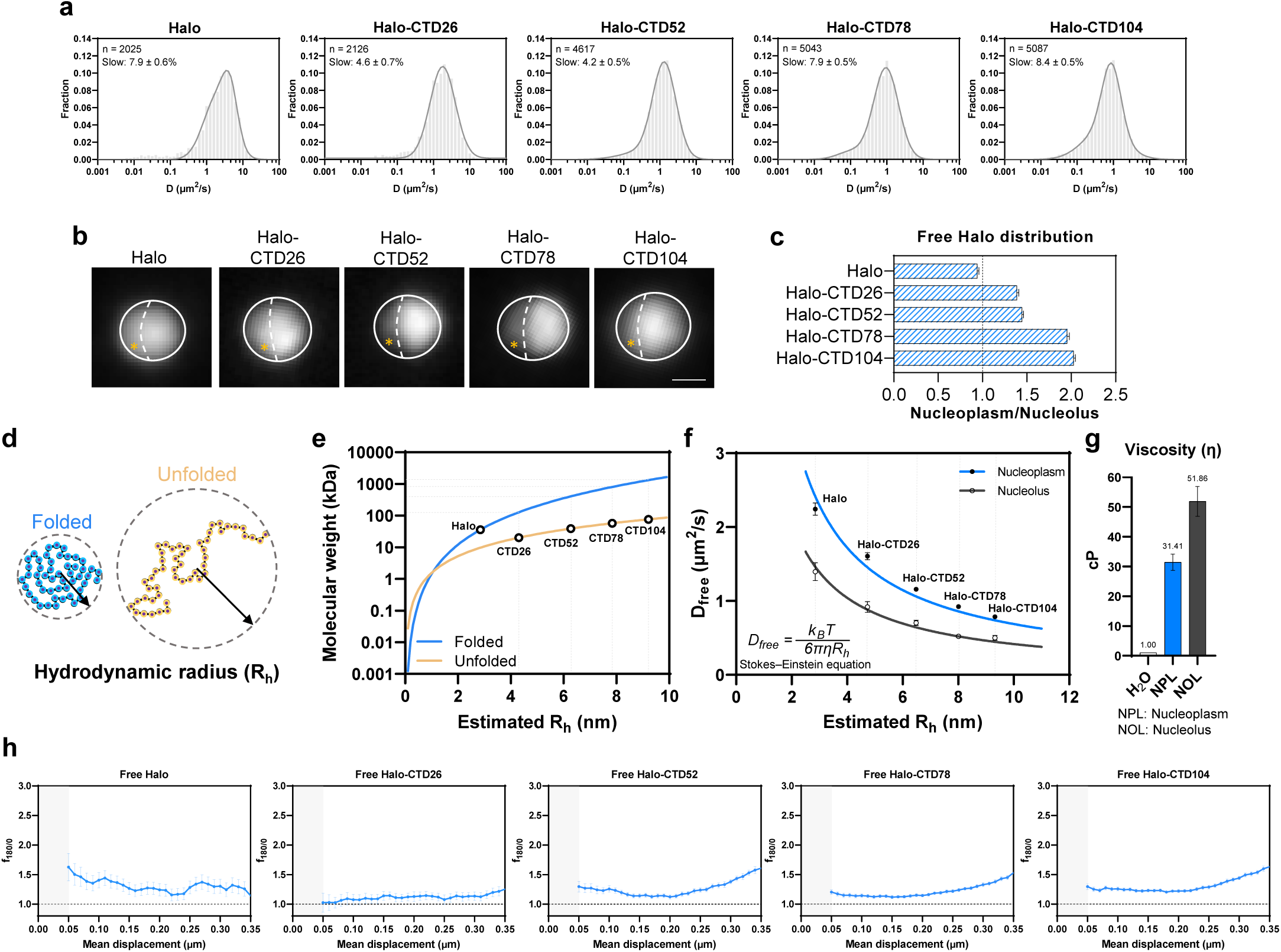
CTD is necessary but not sufficient for confining protein diffusion. **a**, HaloTag (NLSx2-Halo) and Halo-CTD fusions diffusion coefficient histogram by CTD length (n: number of trajectories; mean value ± s.d.). The first plot is reuse of the top panel’s second plot from Extended Data Fig. 1h. **b**, HaloTag and Halo-CTD fusions bulk wide-field staining using JFX650 in live cell. Scale bar: 1.0 µm. **c**, Nucleoplasm/Nucleolus ratios for freely diffusing HaloTag and Halo-CTD fusions (n = 100 resamplings; mean value ± s.d.). **d**, Protein disorder affects hydrodynamic radius (R_h_). **e**, Theoretical relationship between molecule weight and R_h_ for folded and unfolded peptides (Data from Fluidic Analytics). **f**-**g**, **f**, D_free_ and estimated R_h_ of the HaloTag and Halo-CTD fusions in nucleoplasm and nucleolus. Data fitted with Stokes-Einstein equation to calculate **g**, apparent viscosity (η) of yeast nucleoplasm and nucleolus, respectively. K_B_: Boltzmann’s constant. T: Temperature in Kelvin; 295.15K (22°C) (n = 100 resamplings; mean value ± s.d.). **h**, f_180/0_ plot for HaloTag and Halo-CTD fusions (n = 100 resamplings; mean value ± s.d.).

