## Supplementary material for "Disordered C-terminal domain drives spatiotemporal confinement of RNAPII to enhance search for chromatin targets": Methods

### Yeast strain construction

All *Saccharomyces cerevisiae* strains in this study are isogenic derivatives of BY4741 with *pdr5Δ* deletion for enhanced HaloTag ligand labeling. Yeast strains were generated using standard yeast transformation. Free HaloTag was fused with a bipartite nuclear localization signal (NLSx2; KRTADGSEFESPKKKRKV) at the N-terminus for nuclear localization. Strains with *pdr5Δ* and GFP markers exhibit growth identical to the WT (Extended Data Fig. 5i). Strains with defects in promoter scanning and transcription start site (TSS) selection by the PIC exhibit sensitivity to mycophenolic acid (MPA). This sensitivity is reflected in the inability of the PIC to shift to the downstream *IMD2* TSS in the presence of MPA, as observed in certain mutations of TFIIF and TFIIH[63]. Growth of strains with HaloTag fusions to various PIC component subunits was tested on SC-MPA (20 μg/ml) agar plates, and those showing no MPA sensitivity phenotype were selected for further analysis (Extended Data Fig. 5k). The procedure for inserting 24xPP7 into the 5' UTR of *HSP82* was performed as previously described[64]. All $RPB1^{Rpo21}$ CTD length mutants preserve the tip domain. For CTD0 experiments, $RPB1^{Rpo21}$-CTD0-Halo, under native promoter control, was integrated ectopically in $RPB1^{Rpo21}$-CTD26 (WT) AID background (Extended Data Fig. 5e,f,j). This setup ensures cell viability as CTD0 alone is lethal. During imaging, the WT RNAPII-CTD26 was degraded by auxin treatment, thus permitting the investigation of RNAPII-CTD0 dynamics without interference from the WT copy. Strain list is provided in the Supplementary Table 2. Details of CTD mutant constructions are provided in Supplementary Methods and Supplementary Table 1.

### Microscope setup

All imaging was performed using a custom-built Axio Observer Z1 microscope (Zeiss, Germany) equipped with a 150X glycerin immersion objective (ZEISS, Germany) (Supplementary Fig. 1). Data were collected with an EM-CCD camera (C9100-13, Hamamatsu Photonics, Japan) featuring 16 μm physical pixel size and 107 nm pixel size in recorded images. The laser excitation wavelengths used were 555 nm for JF552 and mScarlet-Ix3, 637 nm for JFX650 and miRFP670nano3, and 488 nm for GFP. For simultaneous two-color STORM imaging, emission lights were split by W-VIEW GEMINI (A12801-01, Hamamatsu Photonics, Japan). Detailed information on emission and excitation filters is described previously[65].

### Single-molecule tracking (SMT)

Yeast cultures grown in Synthetic Complete medium (SC) were treated with JF552-HaloTag ligand at the early log phase ($OD_{600}$ = 0.2–0.3) for 4 hours. HaloTag fusions were sparsely labeled with dye concentrations adjusted for factor abundance. Cells were washed at least six times, adhered to concanavalin A-treated coverslips in cell chambers (Invitrogen, Cat. No. A7816) and imaged at room temperature. Initial exposure to a 555 nm laser yielded a bright nuclear signal, followed by transient shelving of fluorophores in a metastable dark state. JF552 spontaneously and stochastically reverted to a fluorescent state, resulting in sparse single-molecule localization[66,67] (Extended Data Fig. 1e). Fast tracking (10 ms/frame) was performed with continuous 555 nm laser irradiation at ~1 kW/cm$^2$ to extract bound and free molecule dynamics (Supplementary Video 2). Static localization error was 18.9 ± 7.7 nm for Halo-H2B in fixed cells, and 22.0 ± 9.0 nm in live cells. Slow tracking (250 ms/frame) was performed with continuous 555 nm laser irradiation at a lower laser power of ~0.035 kW/cm$^2$ to minimize photobleaching and enhance residence time measurement resolution (Supplementary Video 5). We routinely measured and adjusted laser power and alignment throughout imaging sessions to account for minor fluctuations. Imaging sessions lasted two hours, equivalent to one cell cycle in SC at room temperature; Cse4-Halo$^{\text{Internal}}$ was imaged for only one hour due to its short centromeric half-life. Laser exposure during imaging did not appear to induce any significant changes in cell cycle progression (Extended Data Fig. 1f).

### Localizing and tracking single molecules

Single-molecule localization and trajectory linking were conducted using Diatrack software[68]. For fast tracking, a 6-pixel (642 nm) max jump was applied, and a 3-pixel (321 nm) max jump for slow tracking. Only G1 phase cells were analyzed, with nuclear masks manually drawn based on GFP nuclear landmarks to exclude trajectories outside the nucleus (Extended Data Fig. 3a). Single-step disappearances of bound molecules were observed in both fast and slow tracking, indicating accurate identification of single molecule signals (Extended Data Fig. 8a–d)

### Fast-tracking data analysis

*Selection of Nuclei*

A typical imaging session comprises around 40 movies, each containing approximately 3 in-focus cell nuclei. Given the dynamic nature of yeast nuclei, we carefully selected G1 cells exhibiting minimal nuclear drift during imaging (Extended Data Fig. 1g and 3a). This stringent selection process typically resulted in an average of only 5 suitable nuclei per session being chosen for further analysis. For sufficient localization data, all datasets included a minimum of 10 cell nuclei and 1000 trajectories, obtained from at least two independent imaging sessions.

*Multi-parameter classification*

We used vbSPT[69] (adapted script from Hansen et al.[44]) to perform two-state hidden Markov model (HMM) analysis to differentiate transitioning trajectories and separate single-state segments into individual trajectories. We acknowledge the possibility of classifying trajectories into more than two diffusive states. However, Brownian simulations suggest that subdividing a freely diffusing trajectory into multiple states could create artifacts, particularly when the experimental data is dominated by short trajectories (Supplementary Fig. 2a). Short trajectories might lack sufficient information for accurate sub-classification, and any apparent confinement or directionality could stem from partial observation of the inherent randomness (Supplementary Fig. 2).

Following the HMM analysis, we calculated 10 parameters that describe the diffusivity, geometry and angular orientation of each trajectory (Fig. 1a, Extended Data Fig. 2g and Supplementary Methods). We then applied Uniform Manifold Approximation and Projection (UMAP) for dimensionality reduction, which was semi-supervised by the HMM state identified by vbSPT. Lastly, a Gaussian mixture model (GMM) was used to cluster trajectories into free and bound states (Extended Data Fig. 2e,f and Supplementary Fig. 4). The standard deviations for the fraction of free and bound were determined by bootstrapping 100 times.

We compared diffusion coefficients of factors for their identified bound state ($D_{bound}$). Even though elongation factor Paf1 binds to elongating RNAPII, which could theoretically show higher $D_{bound}$ due to 1D movement along the chromatin, our experiments show that its $D_{bound}$ is similar to that of the histones. This suggests the observed $D_{bound}$ in our experimental setup is predominantly influenced by the movement of chromatin itself, rather than by the 1D movement of the factor along it (Extended Data Fig. 1h,i).

*Nucleoplasm/Nucleolus ratio*

To calculate the ratio of freely diffusing molecules between the nucleoplasm and the nucleolus, we counted the number of localizations in both compartments and normalized these values with the respective areas of the nucleoplasm and the nucleolus. Standard deviations were determined by bootstrapping 100 times. We also carried out Brownian simulations to mimic the single-molecule random distribution for a specific nucleoplasm/nucleolus ratio, using custom-written scripts in R.

*Spatiotemporal mapping*

We transformed, normalized, and merged single-molecule localizations from multiple cell nuclei using the centroids of the nucleus and the nucleolus for reference. All yeast nuclei were in G1 phase, displaying distinct nuclear envelope and nucleolar boundaries to ensure that the single-molecule signals were measured on a similar 2D plane. Since there was no landmark to define the angle around the central axis for the 2D projections from multiple cells, mirroring around the central axis was necessary to facilitate visualization[36]. Bins separated by 10 nm were applied across the nuclear area, each with a 150 nm detection radius, and the probability density was calculated based on the number of localizations within this radius (Extended Data Fig. 3b). The choice of the 10 nm bin distance was arbitrarily determined to ensure adequate data point coverage for constructing the heatmap (Extended Data Fig. 3c). Different detection radii were tested, and 150 nm was selected as it provided a suitable resolution for representing density differences across the nucleus (Extended Data Fig. 3c). The density of bound trajectories maps was normalized against total localizations to highlight changes in the bound fraction, whereas density of free trajectories maps was normalized against total free localizations to emphasize confinement. Since the density value does not reflect the actual molecular density, we do not report the specific values. Instead, we present qualitative changes using a continuous spectral scale that transitions from cool colors (blues) for lower densities to warm colors (reds) for higher densities.

For the genome simulation maps, we used 3D genome simulation data from yeast interphase chromosomes[37] (Extended Data Fig. 3d) and projected the genome coordinates onto a 2D plane. Sequence coordinates from the Saccharomyces Genome Database (SGD) were integrated with data on highly expressed genes across multiple gene classes[41,70], as identified in minimal medium[38], to illustrate their probability density distributions on a 2D map (Fig. 1d). We anticipated that the detection of bound trajectories in single-molecule experiments would likely be biased towards genes that are highly expressed; hence, we selected only highly expressed genes from multiple classes for the genome simulation. Similar to the construction of the spatiotemporal map, mirroring around the central axis was conducted. Consequently, genes located on the opposite side along the central axis might appear in close proximity. It is important to note that this density visualization represents the probability density of gene locations within specific classes in defined regions, rather than actual instantaneous gene clustering. Additionally, we present a genome simulation map for total genomic DNA, which mimics the Rabl configuration of yeast chromosomes. It is also essential to acknowledge that the genome simulation map represents only a snapshot of a single thermodynamically stable configuration of the genome and ignores the dynamic nature of chromosomes in the living cell, as well as the variation between individual cells.

*Anisotropy ($f_{180/0}$) analysis*

We isolated the free trajectories in the nucleoplasm by multi-parameter classification, and extracted the angles between two displacements for analysis. We subsampled the trajectories to obtain angles for a total of 30 time intervals. Next, we identified anisotropy by comparing the change of $f_{180/0}$ (fraction of 180° ± 30° over 0° ± 30°) across various length scales of mean displacement (the mean of the two displacements generating the angle)[44]. We restricted our examination to mean displacements of 350 nm or less, given the small size of yeast nuclei and the apparent anisotropy for longer displacements due to nuclear confinement (Extended Data Fig. 6a,b). Standard deviations were determined by bootstrapping 100 times. Misclassifying bound segments as free displacements or having state-transitioning trajectories could significantly increase the $f_{180/0}$ value[44]. We found that our multi-parameter approach outperformed the original vbSPT approach[44] in classifying free and bound displacements for our yeast data (Extended Data Fig. 2b,d,f). In scenarios with increased bound fraction, misclassifying bound displacements as free would increase the anisotropy observed in the free population. In the *ssn3Δ* experiment, our data show an increased RNAPII bound fraction without a corresponding increase in anisotropy (Extended Data Fig. 7h,i), validating our classification for free and bound displacements.

### Slow-tracking data analysis

*Measurement of residence time*

In contrast to fast tracking, we accounted for blinking or transient defocalization of bound molecules by allowing gaps of up to one frame between two localizations. We linked them as one trajectory if they were less than three pixels ($r_{max}$ = 321 nm) apart. This is the upper threshold for the displacements between two successive frames, and 98% of the bound RNAPII, H2B, H3 and H2A.Z displacements were below this threshold (Extended Data Fig. 8e). A survival curve (1-CDF) against time was plotted and fitted with a double exponential decay model:

$$P(t) = f_{sb}e^{-k_{sb}t} + f_{tb}e^{-k_{tb}t}$$

Here, $k_{sb}$ and $k_{tb}$ represent dissociation rates for stable- and transient-binding events, respectively, and $1 = f_{sb} + f_{tb}$ for the fraction of the two components. We do not interpret transient binding as binding to non-specific sites, but rather as a neutral description of the binding duration. The residence times observed are typically underestimated due to factors such as chromatin movements and photobleaching. We use the dissociation rates of stably bound H2B as a control to correct apparent dissociation rates[47] (Extended Data Fig. 8f). Given that fluctuations in the microscope setup could result in variable photobleaching rates, we routinely image

H2B and the protein of interest on the same day[47,71]. To further overcome technical fluctuations, we spike the imaging culture with Halo-H2B cells harboring a labeled nuclear marker Pus1-miRFP670nano3, and co-image the two strains in the same field of view (Fig. 5a).

The corrected decay constant was obtained as $k_{corrected} = k_{observed} - k_{sb,H2B}$, and the corrected residence time is its inverse. Standard deviations were determined by bootstrapping 10,000 times. The stably bound fraction was calculated by multiplying the fraction bound from fast tracking with the fraction of stably bound ($f_{sb}$) in slow tracking. For CTD9 experiments, we used Halo-H2B cells in CTD9 background for correction. The values of $k_{sb,H2B}$ between CTD9 and WT are comparable (Extended Data Fig. 8g,h).

*Measurement of search time*

Search time is the average duration for a molecule to move between stably binding sites (Fig. 5f), accounting for the time and fraction spent in stably bound, transient bound, and free states, as inferred from fast and slow tracking data. Detailed calculations and explanations can be found in previous study[65]. A limitation of this approach is the difference in exposure time used in fast and slow tracking (10 ms/frame versus 250 ms/frame), as slow tracking may not capture exceedingly transient binding events due to its long frame rate. Nevertheless, the estimated search time remains a useful metric for evaluating changes in search kinetics in different mutants and experimental conditions.

### Stochastic optical reconstruction microscopy (STORM)

Yeast expressing HaloTag fusion proteins were labeled with equal concentrations of JF552 and JFX650, incubated for 4 hours, and either fixed in 4% PFA or imaged as live cells. Image acquisition was conducted using ~1 kW/cm$^2$ of 555 nm and 637 nm laser power, 80 ms/frame, and lasted 5-10 minutes for fixed cells or 5 minutes for live cells. W-VIEW GEMINI enabled simultaneous two-color imaging. Channels were manually aligned, and data were corrected for drift and chromatic aberration using the DoM plugin in ImageJ[72]. For fixed cell experiments, fluorophore blinking was removed by pairwise distance distribution correction (DDC)[73].

### HaloTag bulk staining

Yeast expressing HaloTag fusion proteins were labelled with 100 nM of JFX650 in live cell for 4 hours. Image acquisition was conducted using a 637 nm laser at low power. Z-stacks of wide-field images, each consisting of nine slices with a z-interval of 0.3 μm, were captured. Intensity was calculated using average intensity projection. Beam intensity across the field of view was recorded using a fluorescence slide to rectify distortions caused by Gaussian-shaped beam illumination.

### Auxin-induced degradation (AID)

Auxin-induced degradation (AID)[74] of endogenous RPB1$^{Rpo21}$ or MED14$^{Rgr1}$ was achieved by tagging the target protein with AID*-miRFP670nano3 in strain expressing F-box proteins OsTIR1 and AtAFB2 (Extended Data Fig. 5j). Imaging culture was pre-incubated with auxin (I5148; MilliporeSigma, Germany) at a final concentration of 0.5 mM for an hour, and culture was maintained in auxin-medium throughout imaging. Degradation of target protein was confirmed by the depletion of miRFP670nano3 fluorescence signal in living cells (Fig. 3h and Extended Data Fig. 5e,f). To minimize the potential photodestruction of auxin and the generation of related toxic derivatives by strong laser exposure[75], the medium was replaced with fresh auxin medium every 10 minutes during imaging. For the '−Auxin' control, DMSO was used as a substitute for auxin. To further rule out the potential side-effect of auxin, we added auxin to cells expressing ectopic $RPB1^{Rpo21}$-Halo in an $RPB1^{Rpo21}$ AID background. The changes in the bound fraction and $f_{180/0}$ of RNAPII were minimal compared to those in the WT (Extended Data Fig. 5g,h).

### Heat shock

Cells were subjected to heat shock by washing with 39°C SC medium twice in the imaging chamber, then immediately transferring to a preheated 39°C stage-top incubation chamber (Okolab) for imaging.

### Measuring transcription bursting of *HSP82*

Live-cell transcription imaging was performed on 24xPP7-*HSP82* yeast strains expressing a fusion protein consisting of mScarlet-Ix3 and a non-aggregating ΔFG mutant[76] of the dimerized PP7 coat protein (PP7CPx2-mScarlet-Ix3). Z-stacks of wide-field images, each consisting of nine slices with a z-interval of 0.5 μm, were captured every 20 seconds for 40 minutes, using 150 ms exposure with a 555 nm laser at ~0.005 $kW/cm^2$. At least four replicates were acquired per condition, with a minimum of 100 cells showing transcription site signals in total.

Maximum intensity projected images were drift-corrected, and nuclei were manually inspected for presence of transcription sites. Cells undergoing division were excluded from the analysis. Single-frame bursts were removed, and bursts separated by just one frame were merged[77]. To calculate average ON and OFF times, the cumulative distribution functions (CDFs) of the respective time distributions were fitted with gamma and Weibull distributions, respectively[78]. Standard deviations were determined by bootstrapping 10,000 times.

### Protein extraction and Western blotting

Protein extraction was performed using the LiAc/NaOH method[79], followed by SDS-PAGE separation. Proteins were transferred to a PVDF membrane (Bio-rad, USA) using a TE 70 PWR Semi-dry Transfer Unit (GE Healthcare, USA). Blots were blocked with 5% nonfat milk in TBST (Tris-buffered saline with 0.05% Tween 20) for 1 hour at room temperature, incubated with anti-CTD antibody (8WG16; MilliporeSigma, Germany) in 5% nonfat milk in TBST at 4°C overnight, and then with anti-mouse IgG-HRP (NA931; GE Healthcare, USA) in TBST for 1 hour at room temperature. Blots were developed using ECL Prime Western blotting detection reagent (GE Healthcare, USA) and detected on an ImageQuant™ LAS 4000 biomolecular imager (GE Healthcare, USA).

### Statistics

Statistical analyses were performed using GraphPad Prism (version 9.4.0) or RStudio (version 2021.09.1+372; R (version 4.1.2)). Values are mean ± s.d. unless otherwise stated. Significance levels: * $P \leq 0.05$, ** $P \leq 0.01$ and *** $P \leq 0.001$; ns, not significant. For reliable fitting and statistical evaluation, data obtained from at least two independent imaging sessions were resampled by bootstrapping (100 times for fast tracking, and 10,000 times for slow tracking).

### Data availability

Raw single-molecule trajectory coordinates are accessible at Mendeley Data (https://data.mendeley.com/preview/8zkkrwy9bx?a=2bb1642e-d822-4552-b2c1-9a51c0737cc6). Original videos can be provided upon request.

### Code availability

Custom code can be found at GitHub (https://github.com/yhinling/Ling_et_al_2023).
