## Supplementary information for "Disordered C-terminal domain drives spatiotemporal confinement of RNAPII to enhance search for chromatin targets"

#### Supplementary Methods

##### Multi-parameter classification

Single-molecule tracking experiments often yield transitioning trajectories, representing a mixture of multiple diffusive states. This presents a significant challenge in analyzing many chromatin-binding proteins in yeast, as their residence times are relatively short (approximately 5 seconds for chromatin remodelers[1] and PIC components[2]) and the observed trajectories often contain a mix of bound and free segments, with up to 30% of the total trajectories being state-transitioning trajectories for the chromatin remodeler ISW2[1]. Consequently, the overall diffusion characteristics become an average of free and bound states, which may lead to the perception of apparent intermediate populations displaying confinement properties such as small radii of confinement, small diffusion coefficients and high anisotropy. These characteristics could perhaps account for the intermediate populations previously reported for TBP, TFIIA, TFIIB and TFIIE[2], which are not observed here (Fig. 6a) when transitioning trajectories are also taken into account.

Precise state determination is difficult for live cell experiments as trajectories obtained are typically short (Supplementary Fig. 2a). vbSPT employs a variational Bayesian hidden Markov model (HMM) to separate transitioning trajectories into multiple states according to displacement length distributions. The vbSPT algorithm uses the HMM to infer the most probable underlying state transitions, revealing information about the dynamic properties of the tracked particles[3]. However, unsupervised vbSPT can often overfit experimental data[4,5] (Extended Data Fig. 2a). The common approach, which we follow, is to conservatively categorize trajectories into two principal states: free and bound. This is supported by the diffusion coefficient histograms of chromatin-binding proteins, which typically exhibit two Gaussian peaks (Fig. 1b and Extended Data Fig. 1h).

In addition to the two principal states (free and bound), it might be tempting to further dissect the free state into fast (directed) and slow (confined) states, with the slow state potentially indicating entrapment or confinement. However, free trajectories can exhibit both fast and slow behavior due to the partial observation of inherent randomness, as demonstrated by Brownian simulation (Supplementary Fig. 2). Dissecting these free trajectories into specific states becomes more complex when a molecule does not consistently remain in one specific state for sufficient duration. Therefore, we typically adhere to a 2-state model, i.e. free and bound, unless biological evidence justifies differentiating multiple freely diffusive states or when more than two Gaussian peaks are evident in the diffusion coefficient histogram (after careful consideration of the transitioning trajectories).

Population-based analyses such as Spot-On[6] and saSPT[7] allow the evaluation of trajectory diffusion coefficient distribution and the inference of overall population dynamics, while also compensating for defocalization and localization errors. However, these methods tend to overlook state transition, require data fitting to predefined diffusion models, and are unsuitable for spatiotemporal mapping, which requires the classification of individual tracks into specific states. To address these limitations, we have developed a multi-parameter classification pipeline using a track-based approach. The initial step involves applying a two-state hidden Markov model (HMM) to the single-molecule tracking data, effectively segregating the free and bound segments of all transitioning trajectories into distinct single-state trajectories (Extended Data Fig. 2b). Subsequently, we select 10 parameters, informed by literature[8-10] and introduce a novel parameter, 'Angle Bias', to characterize each trajectory with respect to diffusivity, geometry and angular orientation. An advantage of these parameters is their flexibility in fitting the trajectories' properties without requiring a predefined diffusion model. Similar track-based methods have also recently emerged in the field, such as pEMv2[11], Bound2Learn[12] and diffusional fingerprinting[9].

*10 parameters for multi-parameter classification:*

1. **Diffusion coefficient (D)**: A measure of the molecular movement rate through a medium, calculated from the mean squared displacement (MSD) as a function of time:

   $$D = MSD(\tau) \,/\, (4\tau)$$

   $\tau$ is the time lag/interval. MSD is the average squared displacement a molecule moves from its initial position within a trajectory after a given $\tau$. As molecular diffusion in a complex biological system often deviates from simple Brownian diffusion, the diffusion coefficient was derived from the linear part of the MSD curve and should be considered as an 'apparent' diffusion coefficient.

2. **Alpha (α)**: Represents anomalous diffusion behavior in the trajectory, determined from the MSD scaling exponent as a function of time:

   $$MSD(\tau) \propto \tau^{\alpha}$$

   $\tau$ is the time lag/interval. $\alpha$ describes the deviation from normal diffusion, where $\alpha = 1$ represents normal diffusion, $\alpha < 1$ indicates subdiffusion, and $\alpha > 1$ signifies superdiffusion (Supplementary Fig. 2b).

3. **Maximum displacement**: The largest Euclidean distance a molecule moves between any two consecutive steps within a trajectory.

4. **Minimum displacement**: The smallest Euclidean distance a molecule moves between any two consecutive steps within a trajectory.

5. **Mean displacement**: The average Euclidean distance a molecule moves between any two consecutive steps within a trajectory.

6. **Span**: The Euclidean distance between the initial and final positions of a molecule in a trajectory.

7. **Angle Bias**: A measure of the preferential directionality of a molecule's movement, calculated by a weighted average of the angles (θ) between consecutive displacements in the trajectory, with 0° ≤ θ ≤

180°. Each angle is transformed into a score using the equation based on a bipolar sigmoid function:

$$Score = \frac{1 - \frac{2}{1 + e^{\frac{90-\theta}{weight}}}}{1 - \frac{2}{1 + e^{\frac{90}{weight}}}}$$

Weighted averaging emphasizes extreme angles, with the weight set to 20. Angle Bias value is 0 for normal diffusion, > 0 for forward bias, and < 0 for backward bias.

8. **Asymmetry 1**: A parameter quantifying the deviation from a symmetrical shape, typically represented by an ellipse of gyration[10]. Asymmetry 1 is computed using the radius of gyration tensor (T) for a 2D random walk. The principal radii of gyration (R1 and R2) are the eigenvalues of tensor T, where R1 corresponds to the larger principal radius and R2 to the smaller principal radius of gyration.

$$Asymmetry\ 1 = \frac{(R1 - R2)^2}{(R1 + R2)^2}$$

The ratio ranges from 0 to 1, with 0 for circularly symmetric trajectories and 1 for linear trajectories.

9. **Asymmetry 2**: Orthogonal to Asymmetry 1[10],

$$Asymmetry\ 2 = \frac{R2}{R1}$$

The ratio ranges from 0 to 1, with 1 for circularly symmetric trajectories and 0 for linear trajectories.

10. **Aspect ratio**: The ratio between the long and short side of the minimum bounding rectangle. It is 1 for symmetric trajectories and becomes larger for elongated trajectories.

After calculating the 10 parameters for each trajectory, we apply Uniform Manifold Approximation and Projection (UMAP) for dimensionality reduction. This process is intended to separate the trajectory data into two distinct clusters, corresponding to the free and bound states. However, the UMAP projection may not always clearly differentiate these clusters. To refine this categorization, we enhance the separation of the two clusters by semi-supervision using the hidden Markov model (HMM) states identified by vbSPT (Extended Data Fig. 2c–f). Subsequently, we define the clusters using a Gaussian Mixture Model (GMM), and each trajectory is classified into one of two classes: class 1, representing the bound state, or class 2, representing the free state (Extended Data Fig. 2e–g).

**Generation of CTD mutants**

HaloTag fusions and CTD manipulation of *RPO21* involve integrating the *URA3* gene downstream of *RPO21* (*RPO21-URA3*) and replacing *RPO21-URA3* with *RPO21*-CTD-Halo constructs via counter-selection in 5-fluoroorotic acid (5-FOA). All CTD mutants retain the tip domain, and non-consensus heptads are preserved in CTD truncation mutants (Fig. 2a and Supplementary Table 1). For CTD expansion mutants, multiple consensus heptad ‘YSPTSPS’ are inserted between heptads 15 and 16. To facilitate CTD modification, we engineered a CTD26-encoding DNA sequence with silent mutations to create restriction enzyme sites, referred to as nCTD26 (Supplementary Table 1). Yeast strains with nCTD26 exhibit growth rates identical to WT (Extended Data Fig. 5i).

**CTD26 (WT)**

1. FSPTSPT
2. YSPTSPA
3. YSPTSPS
4. YSPTSPS
5. YSPTSPS
6. YSPTSPS
7. YSPTSPS
8. YSPTSPS
9. YSPTSPS
10. YSPTSPS
11. YSPTSPS
12. YSPTSPS
13. YSPTSPS
14. YSPTSPS
15. YSPTSPS
16. YSPTSPS
17. YSPTSPA
18. YSPTSPS
19. YSPTSPS
20. YSPTSPS
21. YSPTSPS
22. YSPTSPN
23. YSPTSPS
24. YSPTSPG
25. YSPGSPA
26. YSPKQDE

QKHNENENSR

**CTD0**

QKHNENENSR

**CTD8**

1. FSPTSPT
2. YSPTSPA
3. YSPTSPS
4. YSPTSPS
5. YSPTSPS
6. YSPTSPS
7. YSPTSPS
8. YSPTSPS

QKHNENENSR

**CTD9**

1. FSPTSPT
2. YSPTSPA
3. YSPTSPS
4. YSPTSPS
5. YSPTSPS
6. YSPTSPS
7. YSPTSPS
8. YSPTSPS
9. YSPTSPS

QKHNENENSR

**CTD10**

1. FSPTSPT
2. YSPTSPA
3. YSPTSPS
4. YSPTSPS
5. YSPTSPS
6. YSPTSPS
7. YSPTSPS
8. YSPTSPS
9. YSPTSPS
10. YSPTSPS

QKHNENENSR

**CTD15**

1. FSPTSPT
2. YSPTSPA
3. YSPTSPS
4. YSPTSPS
5. YSPTSPS
6. YSPTSPS
7. YSPTSPS
8. YSPTSPS
9. YSPTSPS
10. YSPTSPS
11. YSPTSPS
12. YSPTSPS
13. YSPTSPS
14. YSPTSPS
15. YSPTSPS

QKHNENENSR

**CTD20**

1. FSPTSPT
2. YSPTSPA
3. YSPTSPS
4. YSPTSPS
5. YSPTSPS
6. YSPTSPS
7. YSPTSPS
8. YSPTSPS
9. YSPTSPS
10. YSPTSPS
11. YSPTSPS
12. YSPTSPS
13. YSPTSPS
14. YSPTSPS
15. YSPTSPS
16. YSPTSPS
17. YSPTSPA
18. YSPTSPS
19. YSPTSPS
20. YSPTSPS

QKHNENENSR

### nCTD26

```
     TspMI
4618 CCCGGGTTTTCTCCAACTTCCCCAACATACTCTCCTACCTCTCCAGCGTACTCACCAACATCACCATCGT 4687
4688 ACTCACCAACATCACCATCGTACTCGCCAACATCACCATCGTACTCACCTACATCACCATCGTATTCACC 4757
4758 AACGTCACCATCATATTCGCCAACGTCACCATCATATTCGCCAACGTCGCCATCGTATTCTCCAACGTCA 4827
                                                   BmgBI
4828 CCATCGTATTCGCCAACGTCGCCTTCCTACTCTCCCACGTCGCCAAGCTACAGCCCTACGTCTCCTTCTT 4897
             AatII
4898 ATTCTCCGACGTCTCCATCATACTCTCCTACGTCACCAAGTTACAGCCCAACGTCACCAAGTTACAGCCC 4967
           MreI
4968 AACGTCGCCGGCGTATTCCCCAACATCACCAAGTTATAGTCCTACATCGCCTTCATACTCTCCAACATCA 5037
5038 CCATCCTATTCCCCAACATCACCTTCTTACTCTCCCACCTCTCCAAACTATAGCCCTACTTCACCTTCTT 5107
                                                                 Bpu10I
5108 ACTCCCCAACATCTCCAGGCTACAGCCCAGGATCTCCTGCATATTCTCCTAAGCAAGACGAACAAAAGCA 5177
                   XhoI
5178 TAATGAAAATGAAAACTCGAGATGA 5202
```

**Supplementary Table 1.** Amino acid sequence of CTD truncation mutants (top). Non-consensus CTD heptads are highlighted in yellow, and the tip domain is highlighted in blue. Row numbers indicate the number of heptad repeats. For nCTD26, nucleotides coding for heptad repeats are distinguished by different shades of grey (bottom). Silent mutations are marked in red. Restriction sites are boxed. Numbers represent the nucleotide position of the *RPO21* coding sequence.

| Name | Genotype |
|---|---|
| Parent | MATa his3-Δ1 leu2-Δ1 trp1-Δ63 ura3-52 |
| pdr5Δ | MATa his3-Δ1 leu2-Δ1 trp1-Δ63 ura3-52 pdr5Δ::loxP |
| GGEGp | MATa his3-Δ1 leu2-Δ1 trp1-Δ63 ura3-52 pdr5Δ::loxP GAR1-GFP(S65T)-TRP1 ELO3-GFP(S65T)-natMX6 |
| GGEGp-T | GGEGp his3-Δ1::$P_{ADH1}$-OsTIR1-HIS3 |
| GGEGp-A | GGEGp leu2-Δ1::$P_{ADH1}$-AtAFB2-LEU2 |
| GGEGp-TA | GGEGp his3-Δ1::$P_{ADH1}$-OsTIR1-HIS3 leu2-Δ1::$P_{ADH1}$-AtAFB2-LEU2 |
| GGEGp-T, Rpo21-AID* | GGEGp-T RPO21-AID*-miRFP670nano3-hphMX6 |
| GGEGp-A, Rpo21-AID* | GGEGp-A RPO21-AID*-miRFP670nano3-hphMX6 |
| GGEGp-TA, Rpo21-AID* | GGEGp-TA RPO21-AID*-miRFP670nano3-hphMX6 |
| Rpo21-nCTD26 | GGEGp RPO21-nCTD26 |
| Rpo21-CTD0-Halo | GGEGp-TA RPO21-AID*-miRFP670nano3-hphMX6 ura3-52::pRS406-RPO21-nCTD0-3xHA-Halo |
| Rpo21-CTD8-Halo | GGEGp RPO21-nCTD8-3xHA-Halo |
| Rpo21-CTD9-Halo | GGEGp RPO21-nCTD9-3xHA-Halo |
| Rpo21-CTD10-Halo | GGEGp RPO21-nCTD10-3xHA-Halo |
| Rpo21-CTD15-Halo | GGEGp RPO21-nCTD15-3xHA-Halo |
| Rpo21-CTD20-Halo | GGEGp RPO21-nCTD20-3xHA-Halo |
| Rpo21-CTD26-Halo | GGEGp RPO21-nCTD26-3xHA-Halo |
| Rpo21-CTD52-Halo | GGEGp RPO21-nCTD52-3xHA-Halo |
| Rpo21-CTD78-Halo | GGEGp RPO21-nCTD78-3xHA-Halo |
| Rpo21-CTD104-Halo | GGEGp RPO21-nCTD104-3xHA-Halo |
| ssn3Δ Rpo21-CTD26-Halo | GGEGp ssn3Δ::kanMX6 RPO21-nCTD26-3xHA-Halo |
| ssn3Δ Rpo21-CTD9-Halo | GGEGp ssn3Δ::kanMX6 RPO21-nCTD9-3xHA-Halo |
| esc1Δ yku70Δ Rpo21-CTD26-Halo | GGEGp esc1Δ::loxP yku70Δ::loxP RPO21-nCTD26-3xHA-Halo |
| esc1Δ yku70Δ Sir4-Halo | GGEGp esc1Δ::loxP yku70Δ::loxP SIR4-Halo-kanMX6 |
| Rpo21-CTD26 + Rpo21-CTD9-Halo | GGEGp ura3-52::pRS406-RPO21-nCTD9-3xHA-Halo |
| Rpo21-CTD26 + Rpo21-CTD9, Rgr1-Halo | GGEGp ura3-52::pRS406-RPO21-nCTD9 Rgr1-Halo-kanMX6 |
| Rpo21-AID*, Rpo21-Halo | GGEGp-TA RPO21-AID*-miRFP670nano3-hphMX6 ura3-52::pRS406-RPO21-nCTD26-3xHA-Halo |
| Rgr1-AID* Rpo21-CTD26-Halo | GGEGp-TA RGR1-AID*-miRFP670nano3-hphMX6 RPO21-nCTD26-3xHA-Halo |
| Rgr1-AID* Rpo21-CTD9-Halo | GGEGp-TA RGR1-AID*-miRFP670nano3-hphMX6 RPO21-nCTD9-3xHA-Halo |
| Halo-Htb1 | GGEGp Halo-HTB1 |
| Halo-Htb1 Pus1-miRFP670nano3 | GGEGp Halo-HTB1 PUS1-miRFP670nano3-hphMX6 |
| Hsf1-Halo | GGEGp HSF1-Halo-kanMX6 |
| CTD9 Hsf1-Halo | GGEGp RPO21-nCTD9 HSF1-Halo-kanMX6 |
| 24xPP7-HSP82 | GGEGp 24xPP7-HSP82 [$P_{CYC1}$-NLS-PP7CPx2-mScarlet-Ix3-URA3] |
| CTD9 24xPP7-HSP82 | GGEGp RPO21-nCTD9 24xPP7-HSP82 [$P_{CYC1}$-NLS-PP7CPx2-mScarlet-Ix3-URA3] |
| Paf1-Halo | GGEGp PAF1-Halo-kanMX6 |
| Htz1-Halo | GGEGp HTZ1-Halo-kanMX6 |
| Cse4-Halo | GGEGp CSE4-Halo (internal) |
| Sir4-Halo | GGEGp SIR4-Halo |
| Rpa190-Halo | GGEGp RPA190-Halo-kanMX6 |
| Ret1-Halo | GGEGp RE1-Halo-kanMX6 |
| Halo-Hht1 | GGEGp Halo-HHT1 |
| Halo-Spt15 | GGEGp Halo-SPT15 |
| CTD9 Halo-Spt15 | GGEGp RPO21-nCTD9 Halo-SPT15 |
| Taf1-Halo | GGEGp TAF1-Halo-kanMX6 |
| CTD9 Taf1-Halo | GGEGp RPO21-nCTD9 TAF1-Halo-kanMX6 |
| Spt7-Halo | GGEGp SPT7-Halo-kanMX6 |
| CTD9 Spt7-Halo | GGEGp RPO21-nCTD9 SPT7-Halo-kanMX6 |
| Spt8-Halo | GGEGp SPT8-Halo-kanMX6 |
| CTD9 Spt8-Halo | GGEGp RPO21-nCTD9 SPT8-Halo-kanMX6 |
| Rgr1-Halo | GGEGp Rgr1-Halo-kanMX6 |
| CTD9 Rgr1-Halo | GGEGp RPO21-nCTD9 RGR1-Halo-kanMX6 |
| Med1-Halo | GGEGp MED1-Halo-kanMX6 |
| CTD9 Med1-Halo | GGEGp RPO21-nCTD9 MED1-Halo-kanMX6 |
| Halo-Toa1 | GGEGp Halo-TOA1 |
| CTD9 Halo-Toa1 | GGEGp RPO21-nCTD9 Halo-TOA1 |
| Sua7-Halo | GGEGp SUA7-Halo-kanMX6 |
| CTD9 Sua7-Halo | GGEGp RPO21-nCTD9 SUA7-Halo-kanMX6 |
| Tfa1-Halo | GGEGp TFA1-Halo-kanMX6 |
| CTD9 Tfa1-Halo | GGEGp RPO21-nCTD9 TFA1-Halo-kanMX6 |

| | |
|---|---|
| Tfg1-Halo | *GGEGp TFG1-Halo-kanMX6* |
| CTD9 Tfg1-Halo | *GGEGp RPO21-nCTD9 TFG1-Halo-kanMX6* |
| Tfb4-Halo | *GGEGp KIN28-Halo-kanMX6* |
| CTD9 Tfb4-Halo | *GGEGp RPO21-nCTD9 KIN28-Halo-kanMX6* |
| Kin28-Halo | *GGEGp KIN28-Halo-kanMX6* |
| CTD9 Kin28-Halo | *GGEGp RPO21-nCTD9 KIN28-Halo-kanMX6* |
| NLSx2-Halo | *GGEGp ura3-52::pRS406-*$P_{RPO21}$*-NLSx2-Halo* |
| NLSx2-Halo-CTD26 | *GGEGp leu2-Δ1::pRS405-*$P_{RPO21}$*-NLSx2-Halo-nCTD26* |
| NLSx2-Halo-CTD52 | *GGEGp leu2-Δ1::pRS405-*$P_{RPO21}$*-NLSx2-Halo-nCTD52* |
| NLSx2-Halo-CTD78 | *GGEGp leu2-Δ1::pRS405-*$P_{RPO21}$*-NLSx2-Halo-nCTD78* |
| NLSx2-Halo-CTD104 | *GGEGp leu2-Δ1::pRS405-*$P_{RPO21}$*-NLSx2-Halo-nCTD104* |
| Ssl2-Halo | *GGEGp SSL2-Halo-kanMX6* |
| Halo-Ssl2 | *GGEGp Halo-SSL2-Halo* |
| Nut1-Halo | *GGEGp NUT1-Halo-kanMX6* |
| Tfa2-Halo | *GGEGp TFA2-Halo-kanMX6* |
| Tfg2-Halo | *GGEGp TFG2-Halo-kanMX6* |
| Ssl1-Halo | *GGEGp SSL1-Halo-kanMX6* |
| Ccl1-Halo | *GGEGp CCL1-Halo-kanMX6* |
| Tfb3-Halo | *GGEGp TFB3-Halo-kanMX6* |

**Supplementary Table 2.** Yeast strains constructed for this study. For Rpo21-Halo strains, a 3xHA tag is incorporated between the protein linker (GGGGSGGGG) and HaloTag in anticipation of potential pull-down or ChIP experiments. All strains were generated in this study.

**Supplementary Video 1. Single-molecule tracking of RNAPII.**
JF552 signal in red. ER and nucleolar GFP markers in green. For demonstration only, as simultaneous GFP marker imaging affects single-molecule signal-to-noise ratio. Scale bar: 2.0 µm, Recorded frame rate: 30 ms/frame. Playback speed: 30 ms/frame.

**Supplementary Video 2. Fast tracking of RNAPII-CTD26 (WT) and RNAPII-CTD9.**
Nucleolus in grey, nuclear envelope and plasma membrane outlined in solid and dashed white respectively. Scale bar: 1.0 µm. Color scale represents mean instantaneous velocity of trajectory (µm/s). Recorded frame rate: 10 ms/frame. Playback speed: 40 ms/frame.

**Supplementary Video 3. Anisotropic trajectory of free RNAPII.**
Scale bar: 0.2 µm. Recorded frame rate: 10 ms/frame. Playback speed: 50 ms/frame.

**Supplementary Video 4. Sir4-Halo$^{JFX650}$ in WT and *esc1Δ yku70Δ* strain.**
Scale bar: 1.0 µm. Recorded frame rate: 50 ms/frame. Playback speed: 50 ms/frame.

**Supplementary Video 5. Slow tracking kymograph of chromatin-bound RNAPII.**
Recorded frame rate: 250 ms/frame. Playback speed: 100 ms/frame.

**Supplementary Video 6. *HSP82* transcription bursting in WT under non-heat shock condition.**
In addition to strong signal at nuclear transcription site, cytoplasmic mRNAs are visible from 10:40 to 19:20. Scale bar: 1.0 µm. Recorded frame rate: 20 s/frame. Playback speed: 100 ms/frame.

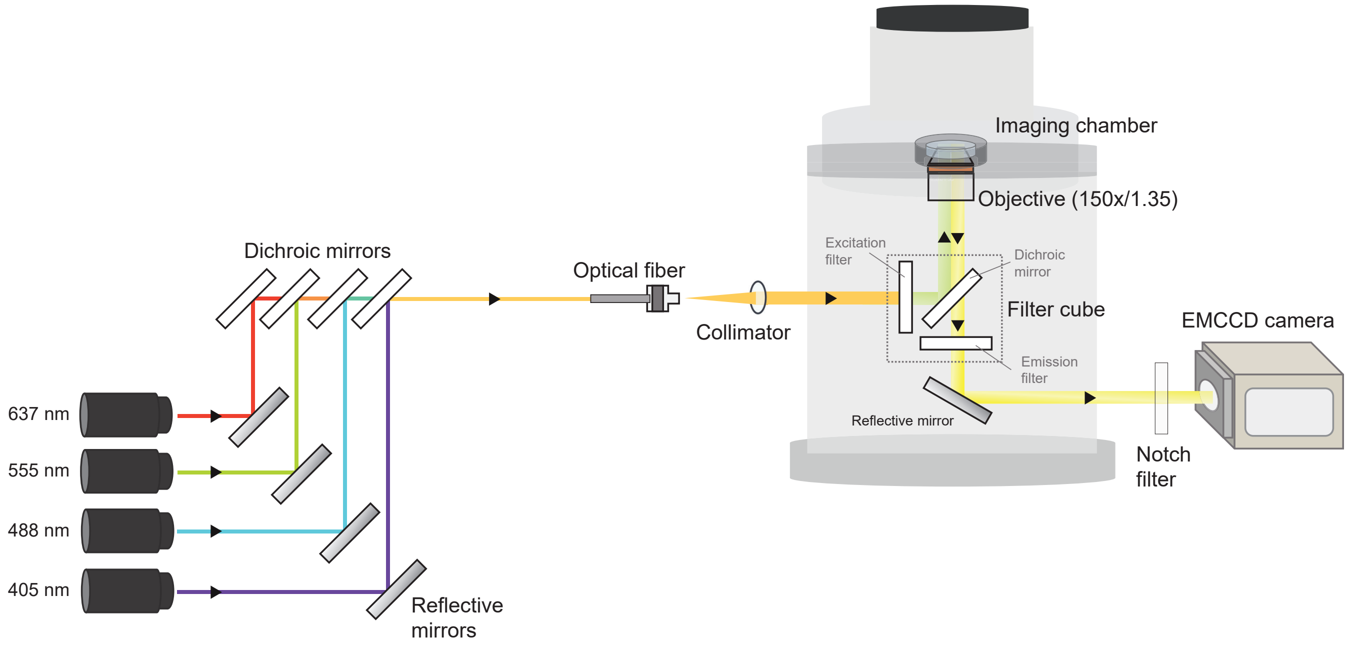


**Supplementary Figure 1. Wide-field microscope setup for single-molecule tracking.**

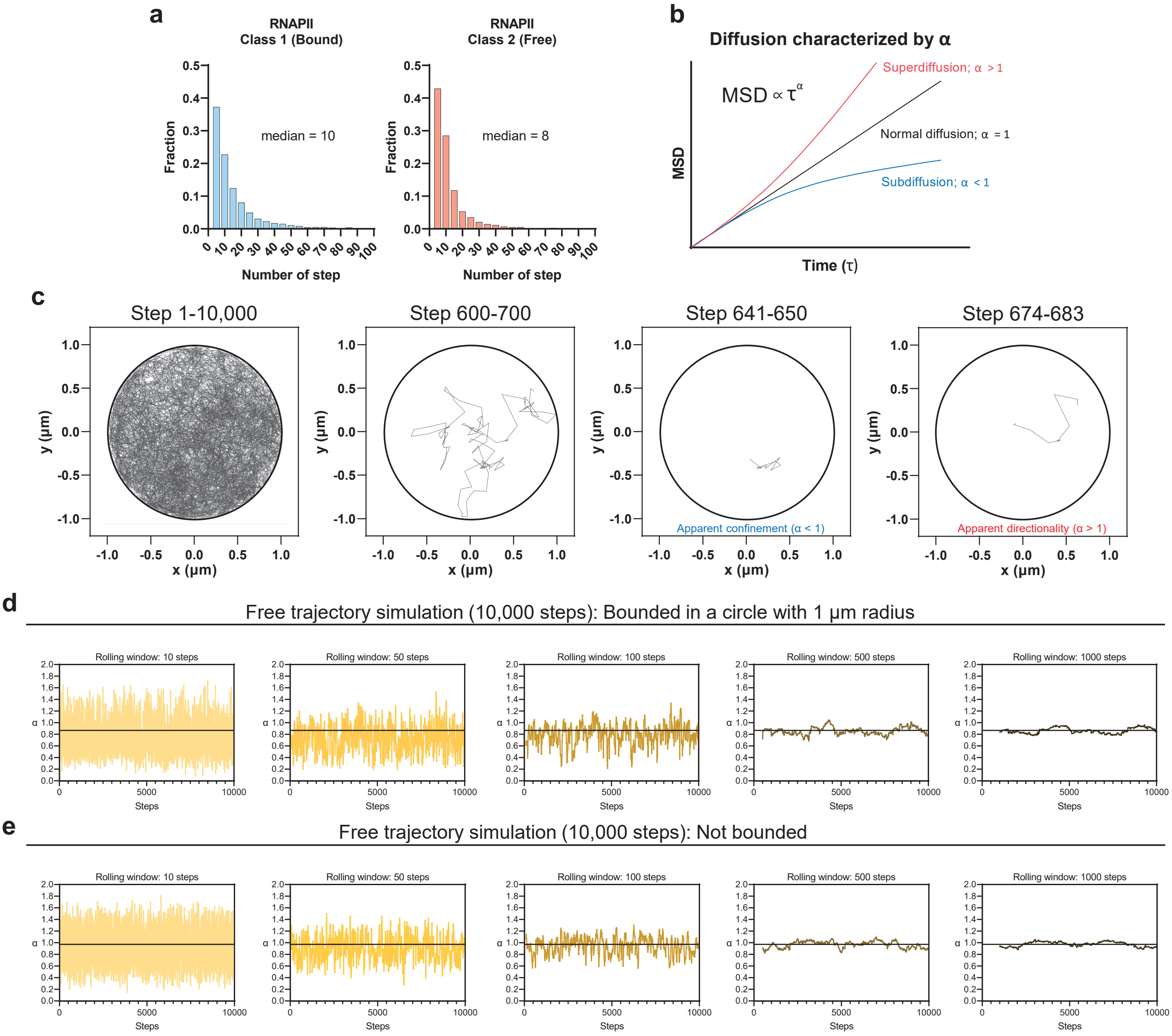


**Supplementary Figure 2. Partial observation of inherent randomness resulted in apparent confinement or directionality.** **a**, Median trajectory length (step) of bound (left) and free (right) RNAPII. **b**, Anomalous diffusion behavior determined by the scaling exponent ($\alpha$) in the mean squared displacement (MSD) as a function of time. **c**, Apparent confinement or directionality for a Brownian simulated trajectory. **d**-**e**, Apparent confinement ($\alpha < 1$) or directionality ($\alpha > 1$) for a Brownian simulated trajectory **d**, bounded within a 1 μm radius circle or **e**, not bounded.

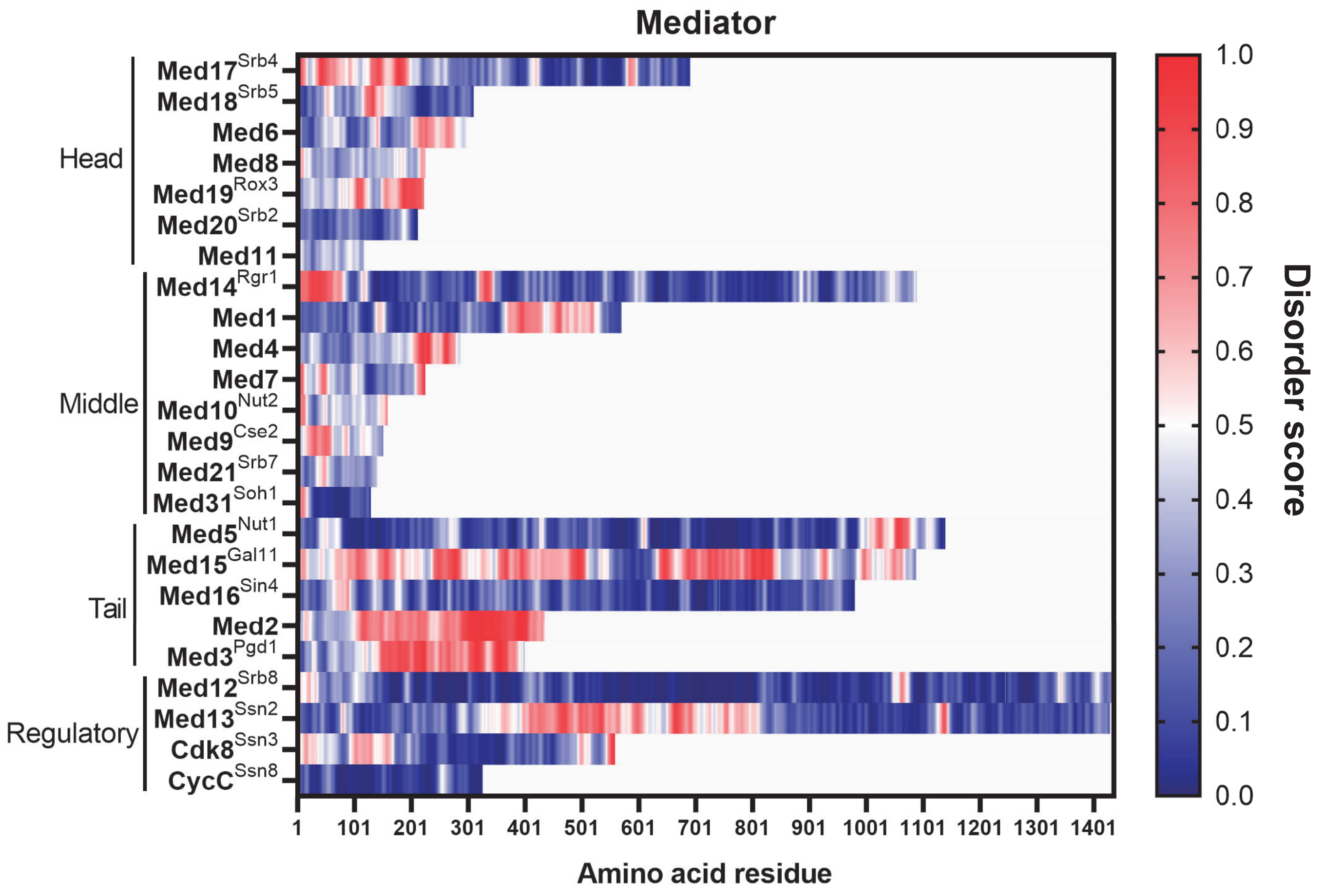


**Supplementary Figure 3. IUPRED3 disorder scores for Mediator subunits.** Residues with predicted scores above 0.5 considered disordered.

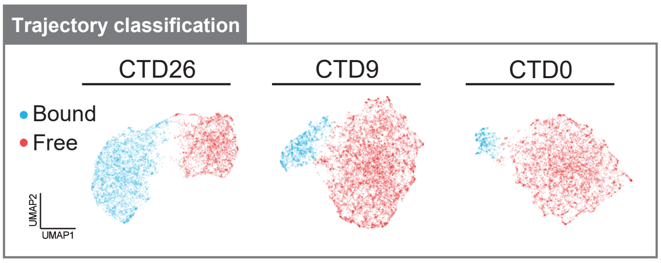


**Supplementary Figure 4. Trajectory classification of RNAPII and CTD truncation mutants.** 10 parameters were calculated from each of the trajectories (Extended Data Fig. 2g and Supplementary Methods) and then subjected to UMAP dimensionality reduction and GMM clustering.
